# Cigarette smoke preferentially induces full length ACE2 exposure in primary human airway cells but does not alter susceptibility to SARS-CoV-2 infection

**DOI:** 10.1101/2021.09.08.459428

**Authors:** Linsey M Porter, Wenrui Guo, Thomas WM Crozier, Edward JD Greenwood, Brian Ortmann, Daniel Kottmann, James A Nathan, Ravindra Mahadeva, Paul J Lehner, Frank McCaughan

**Affiliations:** Department of Medicine, University of Cambridge, Addenbrookes Hospital, Cambridge UK, CB2 OQQ; Cambridge Institute of Therapeutic Immunology & Infectious Disease, Department of Medicine, University of Cambridge, Puddicombe Way, Cambridge, UK, CB2 0AW; Cambridge University Hospitals NHS Foundation Trust, University of Cambridge, Addenbrookes Hospital, Cambridge UK, CB2 OQQ

## Abstract

Cigarette smoking has multiple serious negative health consequences. However, the epidemiological relationship between cigarette smoking and SARS-CoV-2 infection is controversial; and the interaction between cigarette smoking, airway expression of the ACE2 receptor and the susceptibility of airway cells to infection is unclear. We exposed differentiated air-liquid interface cultures derived from primary human airway stem cells to cigarette smoke extract (CSE) and infected them with SARS-CoV-2. We found that CSE increased expression of full-length ACE2 (flACE2) but did not alter the expression of a Type I-interferon sensitive truncated ACE2 that lacks the capacity to bind SARS-CoV-2 or a panel of interferon-sensitive genes. Importantly, exposure to CSE did not increase viral infectivity despite the increase in flACE2. Our data are consistent with epidemiological data suggesting current smokers are not at excess risk of SARS-CoV-2 infection. This does not detract from public health messaging emphasising the excess risk of severe COVID-19 associated with smoking-related cardiopulmonary disease.

## Introduction

SARS-CoV-2 is the causative agent of coronavirus disease 2019 (COVID-19). The SARS-CoV-2 envelope spike (S) protein is essential for virus attachment and cell entry via the main cellular receptor - angiotensin converting enzyme 2 (ACE2) (Kuba et al. 2005; Wang et al. 2020; Daly et al. 2020; Li et al. 2003). Entry is further dependent on S-protein priming by TMPRSS2 facilitating fusion of viral and cellular membranes (Hoffmann et al. 2020; Walls et al. 2020; Shang et al. 2020).

ACE2 is a membrane-associated aminopeptidase expressed in a range of tissues including vascular endothelia, ureteric epithelia and the small intestine (Harmer et al. 2002; Hamming et al. 2004; Zou et al. 2020; Hikmet et al. 2020). In the renin-angiotensin-aldosterone system (RAAS), it converts the vasoconstrictive hormone angiotensin-II to the vasodilator Ang 1-7 but has other physiological roles in glucose homeostasis and beta cell function (Jiang et al. 2014) (Niu et al. 2008; Bindom et al. 2010).

The physiological role for ACE2 at homeostasis in airway epithelial cells is unknown. In preclinical murine models ACE2 was confirmed as the epithelial receptor for SARS-CoV-1 and was shown to confer protection against SARS-CoV-1 associated acute lung injury (Imai et al. 2005; Kuba et al. 2005; Ren et al. 2006). However, the mechanism by which ACE2 mediates protection is unclear and may relate to its role in the pulmonary vascular endothelium rather than the airway epithelium (Imai et al. 2005). In transcriptomic studies of the human respiratory tract and lung, there is a proximal-distal ACE2 mRNA expression gradient; expression is highest in the nasal epithelium and lower distally in the alveolar epithelium, mirroring the permissiveness to SARS-CoV-2 infection (Sungnak et al. 2020; Lukassen et al. 2020; Ziegler et al. 2020; Hou et al. 2020) (Zamorano Cuervo and Grandvaux 2020). The distribution of ACE2 protein expression is less well characterised due to the paucity of validated reagents but is consistent with the proximal-distal graded expression pattern (Ortiz et al. 2020; Hikmet et al. 2020). Notably, ACE2 expression in the lung is not altered by ACE inhibitors or angiotensin receptor blockers (Lee et al. 2020).

In addition, an N-terminally truncated (dACE2) isoform that is sensitive to interferon stimulation or viral infection has been detected (Onabajo et al. 2020; Ng et al. 2020; Blume et al. 2021; Shajahan et al. 2020). Importantly, dACE2 does not express the SARS-CoV-2 spike-protein binding domain and its relevance in SARS-CoV-2 infection and normal physiology remains unclear (Onabajo et al. 2020; Blume et al. 2021).

The impact of ACE2 expression on COVID-19 incidence and severity is unclear (Chung et al. 2020). Much of the available information on ACE2 expression is from scRNA databases and has not reported the relative expression of the two isoforms. Given the differential isoform binding to SARS-CoV-2, this is likely to be an important issue.

Smoking has been associated with increased ACE2 expression in rodent models and in studies on human subjects (Gebel et al. 2010; Hung et al. 2016; Yilin, Yandong, and Faguang 2015; Cai et al. 2020; Brake et al. 2020; Leung et al. 2020; Smith et al. 2020). Consistent with this, higher ACE2 transcripts have been detected in the lungs of COPD patients (Cai et al. 2020; Jacobs et al. 2020; Leung et al. 2020; Smith et al. 2020) and in airway cells from occasional or “social” smokers exposed to 3 cigarettes over a 24hr period (Aliee et al. 2020). Importantly, studies to date have not linked RNASeq data with protein and isoform expression.

The epidemiological data regarding the association of smoking with COVID-19 is conflicting and controversial (Farsalinos et al. 2020; Simons et al. 2020; Rossato et al. 2020; Hopkinson et al. 2021; Grundy et al. 2020; Patanavanich and Glantz 2020). However, an extensive and ‘living’ meta-analysis undergoing regular updates as the evidence improves, suggests that current smoking is not associated with an increased risk of SARS-CoV-2 infection (Simons et al. 2020). Large surveys have suggested that chronic respiratory disease including COPD (mainly a smoking-related disease in the UK) may be associated with an increased risk of severe COVID-19 (Docherty et al. 2020; Williamson et al. 2020). This may be consistent with the observation in the living meta-analysis that former smokers were at an increased risk of severe COVID-19 (Simons et al. 2020; Zhao et al. 2020).

Differentiated human airway epithelial cells grown at the air-liquid-interface (ALI) are the optimal laboratory system in which to model the impact of smoking on the early stages of SARS-CoV-2 infection (Blume et al. 2021; Purkayastha et al. 2020; Sachs, Finkbeiner, and Widdicombe 2003). Purkayastha and colleagues recently reported brief exposure of primary airway cultures (3 minute/day for four days) to “headspace” cigarette smoke (CS), that is CS in a closed environment, to investigate the impact of CS on ACE2 expression and SARS-CoV-2 infection (Purkayastha et al. 2020). A significant increase in ACE2 expression was not detected in response to CS, in contrast to the available molecular epidemiological data detailed above. However, they reported that CS exposure did increase viral infection at 48 hours.

We now report our observations from experiments in which we exposed primary human bronchial epithelial cells (HBECs) at ALI to cigarette smoke extract (CSE). We find that HBECs upregulate ACE2 expression in response to CSE – consistent with the molecular epidemiological evidence (Cai et al. 2020; Brake et al. 2020). However, this did not lead to an increase in infection by SARS-CoV-2 despite preferential upregulation of full-length ACE2 receptor (flACE2) rather than the N-terminal truncated isoform in response to CSE exposure. This suggests that in normal human bronchial epithelial cells physiological expression of flACE2 does not limit viral infectivity. We go on to show that dACE2 is a Type I interferon-sensitive gene in primary HBECs, and define the impact of CSE, nicotine and NRF2 agonists on ACE2 isoform expression.

This study directly addresses one of the controversies in the link between smoking and SARS-CoV-2. Our results are consistent with epidemiological evidence suggesting that current smoking is not associated with a higher incidence of SARS-CoV-2 infection.

## Results

### ACE2 is expressed on differentiated ciliated cells at homeostasis

Previous studies have shown ACE2 expression increases following differentiation at the air-liquid interface (ALI), but could be reversed if cultures were resubmerged (Jia et al. 2005). We grew HBECs from Donor 1 at the air-liquid interface (ALI) for a minimum of 4 weeks to produce a well-differentiated, pseudostratified mucociliary epithelium (**Figure 1a**). ACE2 mRNA expression was increased on differentiation, as was TMPRSS2, the main cellular protease implicated in SARS-CoV-2 spike protein cleavage and FOXJ1, a key transcription factor regulating airway ciliation. Full length ACE2 protein was not detectable in submerged HBEC cultures but was readily detectable on differentiation (**Figure 1b-d**) emphasising the importance of using differentiated HBECs to model airway infection. Confocal imaging demonstrated apical ACE2 expression colocalising with markers of ciliated cells but not goblet cells (**Figure 1 e,f**) (Sims et al. 2005).

**Figure 1.**
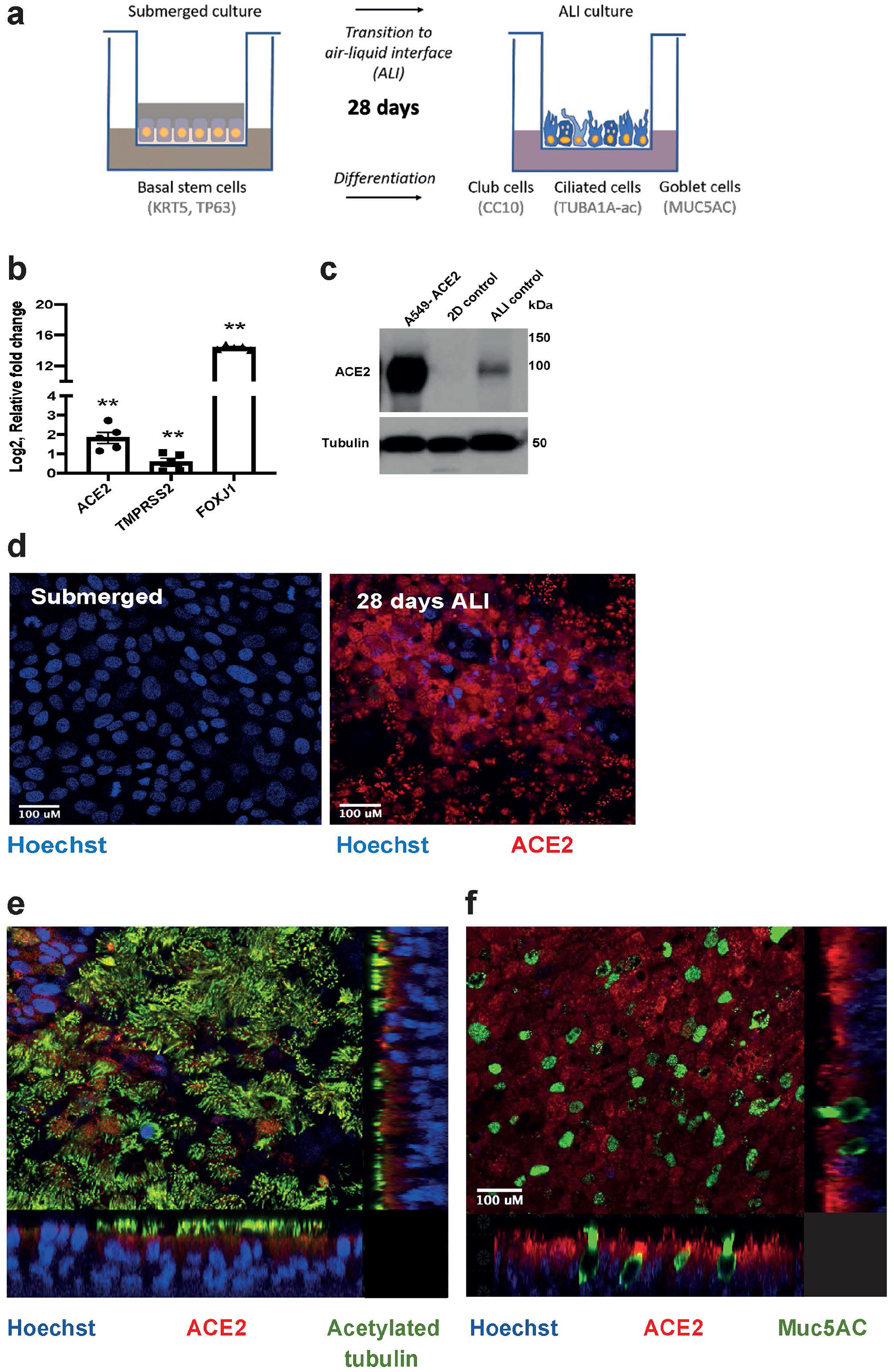
ACE2 expression increases upon differentiation in HBECs cultured at the air-liquid interface (ALI). **A.** Schematic representation of ALI experimental set-up using HBECs. Cell type specific markers are shown in parentheses. **B.** HBEC *TMPRSS2* and *ACE2* expression (RNA) both increase when cultured for 28 days at the ALI compared to submerged, non-differentiated cell culture. Expression of the transcription factor required for ciliation, *FOXJ1* is also significantly upregulated. RT-qPCR data presented as log2 relative fold-change in expression compared to submerged HBECs from n=5 independent experiments (Mann Whitney, **, P < 0.01). Error bars represent mean and the standard error of the mean. **C.** ACE2 protein is also increased during differentiation. A549 cells overexpressing ACE2 are used as a positive control. Representative western blot from 3 independent experiments. ACE2 antibody Ab228349 was used. **D.** ACE2 expression (red fluorescence, antibody Ab228349) is upregulated on differentiation. **E**) ACE2 (red, Antibody used 21115AP) is expressed apically on the epithelial cell surface, predominantly colocalising with ciliated cells (acetylated tubulin, green fluorescence). **F) ACE2** (red, antibody 21115AP) does not colocalise with goblet cells (MUC5AC, green fluorescence). Scale bars on fluorescent images = 100\’fm. All experiments in Figure 1 used cells expanded from Donor 1.

### Cigarette smoke extract increases ACE2 expression in differentiated HBECs

We then exposed differentiated ALI cultures from Donor 1 to 10% cigarette smoke extract (CSE) for 48 h before harvesting cells (**Figure 2a**). CSE exposure induced a significant increase in ACE2 (mRNA) and marked induction at the protein level relative to controls (**Figures 2b** and **2c**). We evaluated ACE2 immunofluorescence after CSE exposure which was consistent with increased apical ACE2 expression relative to control wells (**Supp Figure 1**). Increased ACE2 levels were also detected from differentiated ALI cultures derived from Donor 2, a former smoker (61 pack-years) with COPD (**Supplementary Figure 2**). Of note, there was no evidence of cytotoxicity in response to CSE, as exposure did not cause an increase in apoptosis or necrosis as shown by flow cytometric analysis (**Figure 2d**; **Supplementary Figure 3**) or an obvious cytopathic effect on histology (**Figure 2e**). Importantly, given prior data on the impact of nicotine on ACE2 expression in submerged undifferentiated bronchial epithelial cells, (Russo et al. 2020), we found that nicotine did not significantly alter the expression (mRNA) of either ACE2 or lead to a consistent change in ACE2 protein expression (**Supplementary Figure 4**). CSE had a negligible effect on the expression of the predominant nicotinic acetylcholine receptor expressed on airway epithelial cells – CHRNA7 (**Figure 2f,g**).

**Figure 2.**
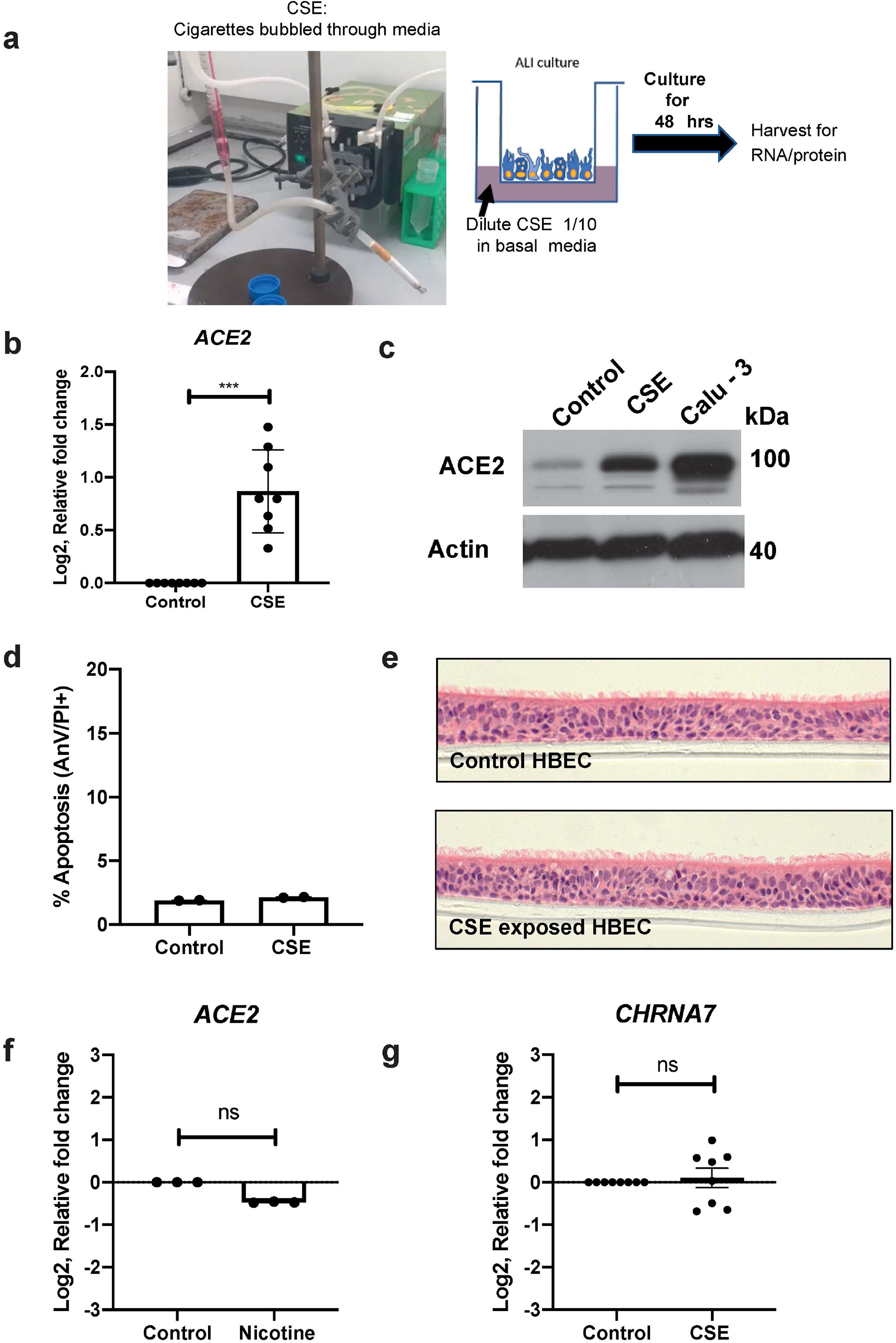
Exposure to cigarette smoke extract (CSE) increases ACE2 expression at both the RNA and protein level. **A.** Schematic showing ALI culture experimental set-up with CSE. **B.** CSE exposure (10%) for 48 h increases *ACE2* expression in differentiated HBECs relative to untreated controls. RT-qPCR data presented as log2 relative fold-change in expression from n=8 independent experiments (Mann-Whitney, ***, P < 0.001). **C.** CSE exposure (10%) for 48 h also increases ACE2 protein expression relative to untreated control. Calu-3 are presented as the positive control. Representative western blot from 3 independent experiments. ACE2 antibody – ab15348. **D.** CSE exposure does not induce apoptosis in differentiated HBECs at ALI relative to control samples as analysed by flow cytometric analysis using AnnexinV (AnV) and Propidium iodide (PI) staining see also Supplementary Figure 3. Data represents 2 independent experiments using cells from Donor 1. **E.** H&E staining of sectioned differentiated ALI HBEC cultures from Donor 1 with and without CSE exposure (x20 magnification). **F**. Treatment of differentiated HBECs at ALI with 1 uM Nicotine for 48 h does not induce *ACE2* expression. **G**. CSE exposure does not significantly alter *CHRNA7* expression **G**. RT-PCR data for **F** and **G** is presented as log2 relative fold-change in expression from n=3-8 independent experiments (Mann-Whitney, ns).

### HBECs exposed to CSE are not more susceptible to infection by SARS-CoV-2

To determine whether CSE exposure would render the cells more susceptible to SARS-CoV-2 infection, differentiated ALI cultures (Donor1) were pre-treated with CSE for 48 h, then inoculated with SARS-CoV2 for 3 h and harvested after 72 hours for flow cytometric quantitation of infection or immunofluorescence (IF) (**Figure 3a**).

**Figure 3.**
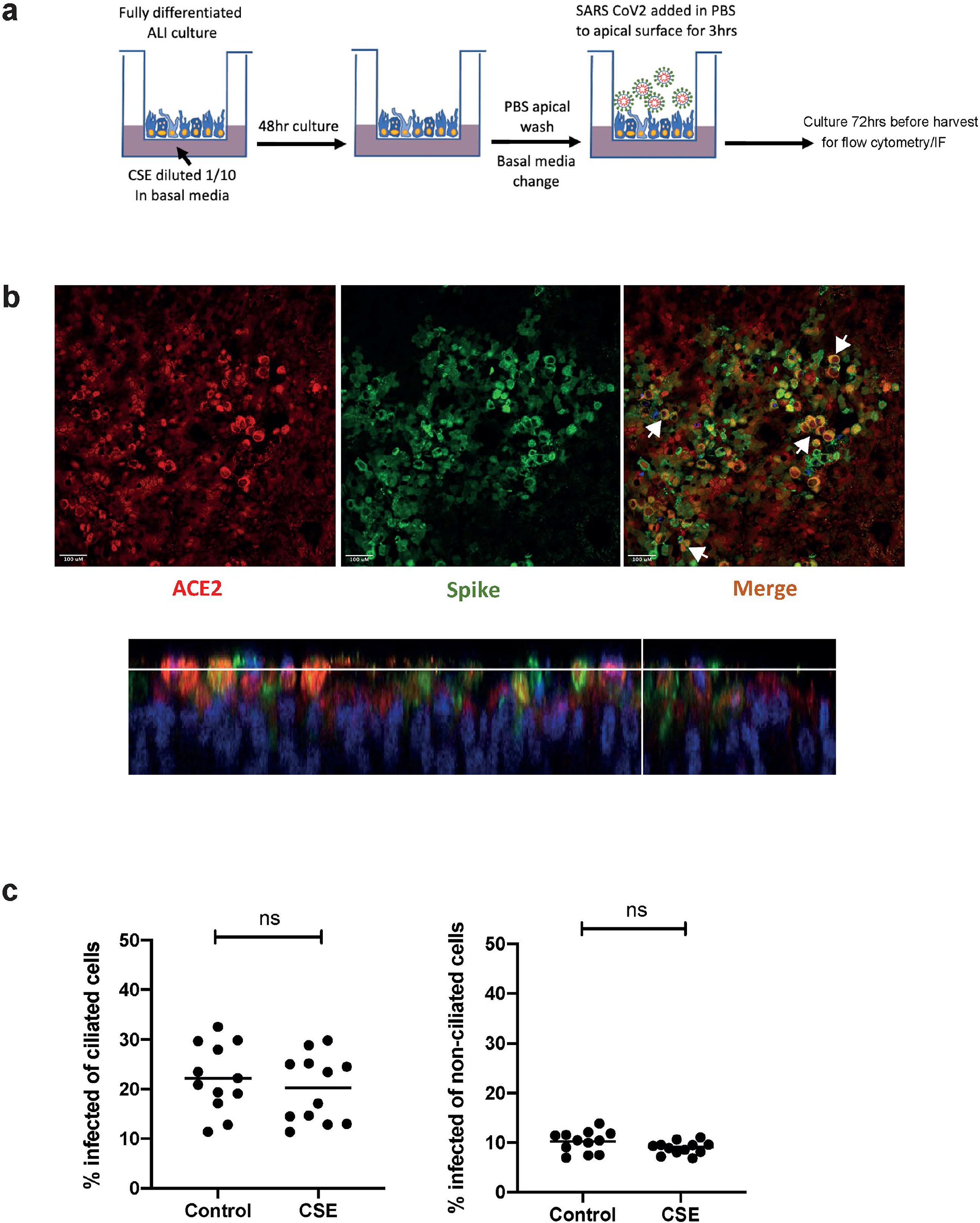
HBECs exposed to CSE are not more susceptible to infection by SARS CoV2. **A.** Schematic representation of SARS-CoV-2 infection of CSE exposed HBEC ALI cultures. **B.** SARS-CoV-2 infection was detected using an antibody specific to the viral spike protein (S2 domain). Representative immunofluorescent images showing 72 h post SARS-CoV-2 infection; expression of viral spike protein (green) primarily co-localised with ACE2 (red) expressing cells. White arrowheads indicate co-localisation of markers in merged imaged. **C.** Flow cytometry quantification of ciliated and non-ciliated cells infected with 8×10^3^ TCID50 of B.29 lineage SARS-CoV-2 following control/CSE exposure (n=12); each dot represents an individual transwell from 2 independent experiments, bars represent mean values (Mann Whitney; ns, non-significant).

ACE2 colocalised with spike protein in infected wells (**Figure 3b**) consistent with ciliated cells being more susceptible to infection and with prior reports (Sims et al. 2005; Schaefer et al. 2020). On some specimens infected cells appeared to be extruded from the epithelial surface – as previously reported for SARS-CoV-1 and 2 (Sims et al. 2005).

Importantly, despite the increased ACE2 levels following CSE exposure, there was no significant difference in the total infected fraction or infected ciliated cell fraction between control or CSE exposed cells (**Figure 3c**). Therefore, in this model CSE exposure did not increase SARS-CoV-2 infection.

### Regulation of ACE2 isoform expression – impact of cigarette smoke

The relationship between cigarette smoke exposure and viral infections has previously been investigated using primary cells – for example cigarette smoke was associated with increased rhinovirus infection in submerged cultures (Eddleston et al. 2011) or influenza A infection in differentiated ALI cultures (Duffney et al. 2018). However, SARS-CoV-2, in contrast to typical respiratory viruses, is associated with an attenuated cellular interferon response (Blanco-Melo et al. 2020). Of note, a truncated isoform of ACE2 denoted as dACE2 was recently implicated as an interferon-sensitive gene (ISG), but importantly predicted to not act as a receptor for SARS-CoV-2 based on the reported ACE2-Spike receptor binding domain interaction (Blume et al. 2021; Onabajo et al. 2020; Ng et al. 2020; Li et al. 2005; Lan et al. 2020).

We therefore explored the impact of cigarette smoke exposure on ACE2 isoform expression. using recently described tools (**Figure 4a**) (Onabajo et al. 2020). We first assessed the relative expression of flACE2 and dACE2 in submerged primary HBECs compared to differentiated HBECs at ALI. dACE2 was upregulated on differentiation but modestly compared to the full-length receptor (**Figure 4b**).

**Figure 4.**
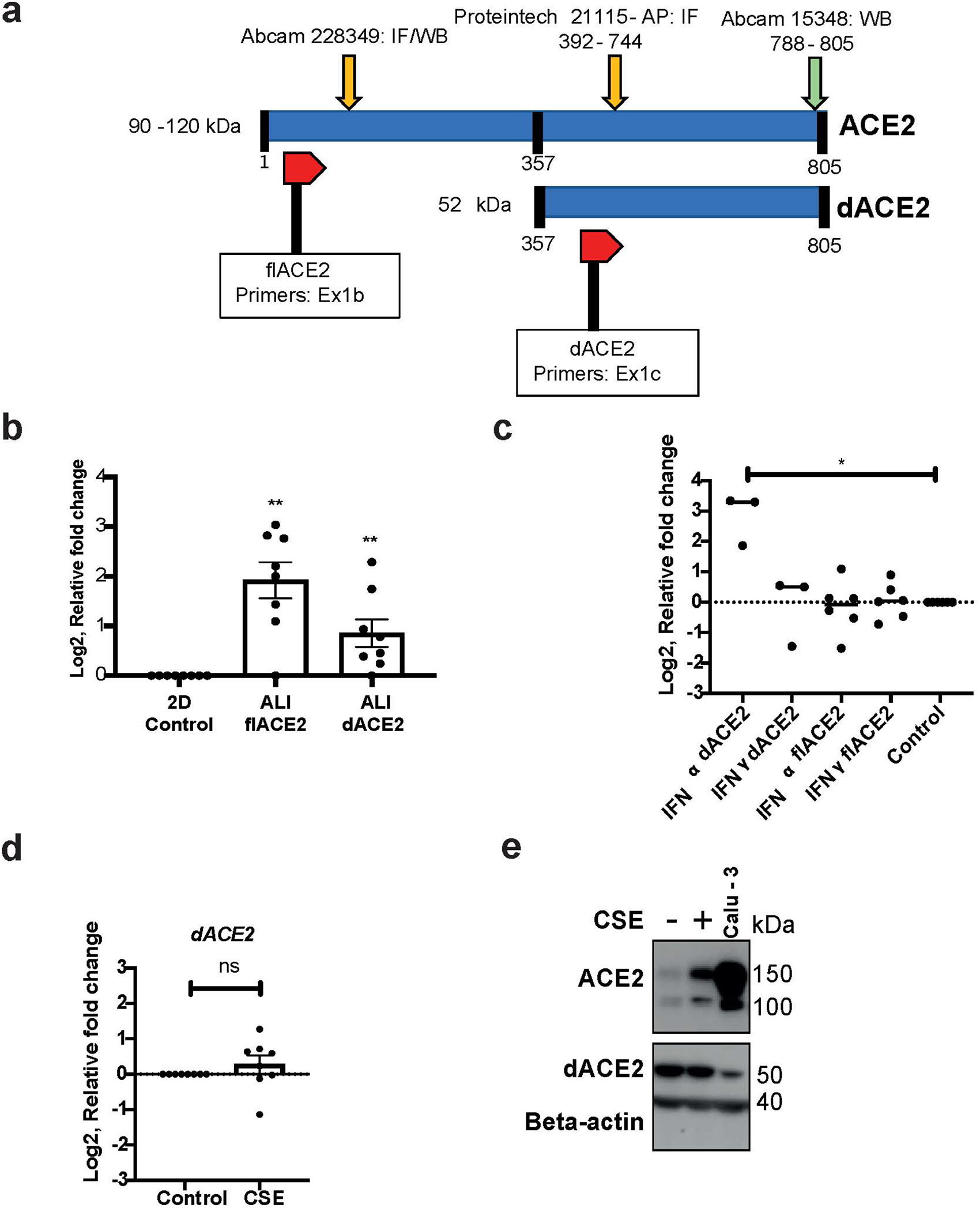
A short isoform of ACE2 is upregulated during HBEC differentiation and interferon-alpha stimulation but not CSE exposure at ALI. **A.** Schematic of full-length ACE2 (flACE2) and the truncated isoform (dACE2) detailing position of antibody binding epitopes for immunofluorescence and western blot analysis. The location of primers used to distinguish ACE2 and dACE are also shown. **B.** HBECs differentiated at ALI upregulate a short isoform of ACE2 (dACE2) as well as the full length ACE2. RT-PCR data shows log2 relative fold-change in expression from n=7 independent experiments (Mann-Whitney **, P < 0.01). **C.** d*ACE2* is specifically sensitive to interferon-alpha stimulation (24 h) but not interferon-gamma at 24 h. Full-length ACE2 shows no modulation with interferon treatment at 24 h. RT-PCR data shows log2 relative fold-change in expression from n=3-6 independent experiments (Mann-Whitney, * p<0.05). **D**. 48 h exposure of CSE does not promote an increase in dACE2 mRNA. RT-PCR data shows log2 relative fold-change in expression from n=7 independent experiments (Mann-Whitney, ns). **E**. Western blot is representative of 5 independent experiments and shows the impact of 48 h exposure of CSE on flACE2/dACE2 expression. Also see Supplementary Figure 6. ACE2 antibody used ab15348.

We next tested isoform-specific expression of ACE2 exposed HBECs to Type I and Type II interferons and CSE. IFN-α (Type I) but not IFN-γ (Type II) led to a transcriptional upregulation of dACE2 (mRNA). Neither interferon significantly altered the expression of the full-length transcript (**Figure 4c**). This extends the recently published data from immortalised airway cells (Blume et al. 2021) and confirms that the N-terminus truncated transcript (dACE2) is an interferon-sensitive isoform. Further, CSE did not significantly alter the expression of a panel of interferon-sensitive genes (**Suppl Figure 5**). Therefore, differentiated airway epithelial cells have the capacity to respond to Type I interferon and CSE does not mimic that response.

**Figure 5.**
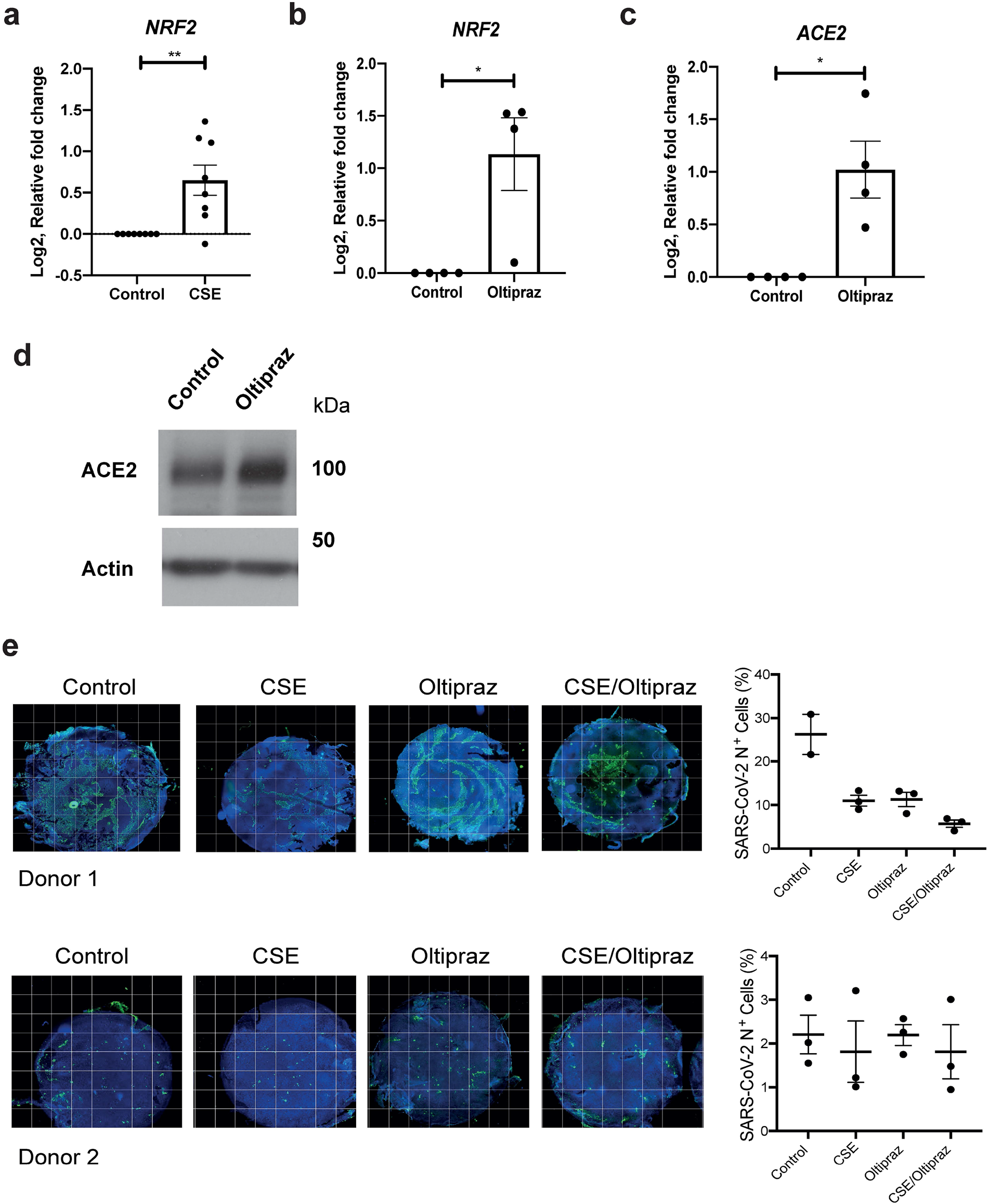
CSE and oltipraz increase ACE2 and NRF2 expression but not SARS-CoV-2 infection. **A.** CSE exposure induces NRF2 mRNA expression at 48 h. **B**. The NRF2 agonist Oltipraz increases NRF2 mRNA **C.** Oltipraz increases flACE2 mRNA expression. D. Oltipraz increases flACE2 protein expression (ACE2 antibody Ab15348). RT-PCR data shows log2 relative fold-change in expression from n=7 independent experiments (Mann-Whitney, ** P < 0.01, * p<0.05). Representative western blot from 3 independent experiments. **E.** Fluorescent image analysis of whole transwell ALI cultures show that for Donors 1 and 2, CSE or NRF2 agonists, alone or in combination, did not result in increased infection. Each datapoint represents a single well and infection was with 1×10^4^ TCID50 of B.1.1.7 SARS-CoV-2. Representative microscopy montages show the entire ALI transwell. Scale bar: 500 μm. Infection was quantitated and presented as percentage of cells infected for each condition.

CSE did not significantly alter the expression of dACE2 mRNA (**Figure 4d**). In agreement with the transcriptional data, both N & C-terminus ACE2 antibodies (**Figure 4a**) showed that CSE consistently upregulated flACE2 protein but had no impact on an ACE2 band migrating at 52kd - the predicted molecular weight of dACE2 (**Figure 4e and Suppl Figure 6**) (Blume et al. 2021). We conclude from these experiments that cigarette smoke does not activate interferon signalling or ISGs in normal human bronchial epithelial cells and preferentially upregulates flACE2 rather than dACE2.

### Antioxidants upregulate ACE2 in differentiated airway epithelial cells

Nuclear factor erythroid 2–related factor 2 (NRF2) is the master transcriptional regulator of the cellular antioxidant response and already a focus of therapeutic efforts to counteract epithelial oxidative stress in COPD. NRF2 agonists have also been proposed as therapeutics for COVID-19 (Olagnier et al. 2020). In our experiments, CSE treatment of ALI cultures led to the expected increase in NRF2 as well as ACE2 upregulation (**Figure 5a**). Further, oltipraz, a KEAP1 inhibitor and NRF2 agonist already in Phase III clinical trials, increased both ACE2 mRNA expression and flACE2 protein (**Figure 5b-d**). This was a consistent finding in two donors – a non-smoker (Donor 1) and an individual with COPD (Donor 2), (**Supplementary Figure 7a**). Despite elevating flACE2 levels (Supplementary Figure 7a), oltipraz pre-treatment did not increase SARS-CoV2 infection (**Figure 5e, Supplementary Figure 7b**). Of note, infection for these experiments was undertaken with the B1.1.7 variant. As others have shown there is considerable inter-experiment variation when using SARS-CoV-2 to infect in primary HBECs (Hou et al. 2020; Ravindra et al. 2021). (However, in three experiments with multiple technical replicates from two donors (**Figure 5e, Supplementary Figure 7b**), there was no increase in infection with either CSE (consistent with **Figure 3C**) or oltipraz. We also assessed the impact of combined treatment with oltipraz and CSE. Again, despite induction of ACE2 (**Supplementary Figure 7a**), there was no increase in SARS CoV-2 infectivity. Therefore, smoking and KEAP1 inhibition both increase flACE2 expression but this does not lead to an increase in SARS-CoV-2 infectivity measure using either flow cytometry or in situ immunofluorescence of differentiated bronchial epithelial cells at the air-liquid interface.

## Discussion

The severity of COVID-19 is associated with increasing age and comorbidities including obesity, diabetes mellitus and chronic respiratory disease. Separately, there is a long established and direct link between smoking and serious adverse health outcomes, including chronic respiratory diseases, cardiovascular disease and cancer.

During the pandemic, there has been intense interest in the link between cigarette smoking and COVID-19. In terms of chronic respiratory disease, smoking is a major global cause of COPD, and current smokers or individuals with COPD are more at risk of severe COVID-19 infections and death (Alqahtani et al. 2020; Leung et al. 2020; Zhao et al. 2020; Guo 2020; Hopkinson et al. 2021; Simons et al. 2020). Molecular epidemiological studies have linked COPD with an increased expression of ACE2, the main receptor for SARS-Co-V2 (Smith et al. 2020; Jacobs et al. 2020; Brake et al. 2020). Further, bulk and single cell RNA-Seq datasets comparing smokers and never-smokers has consistently shown that cigarette smoking (acute and chronic) leads to an increase in ACE2 expression, although these studies have not discriminated between the two ACE2 isoforms that are now known to be expressed in the airway (Cai et al. 2020; Zhang, Yue, et al. 2020; Aliee et al. 2020; Leung et al. 2020).

The elevated ACE2 expression in smokers has been linked with increased susceptibility to infection (Purkayastha et al. 2020; Cai et al. 2020). However, the available clinical epidemiological data suggests that smokers and non-smokers have similar risks of infection, but that those smokers or ex-smokers with cardiorespiratory end-organ damage (COPD, cardiovascular disease) are more likely to have severe infections or die from COVID-19 (Williamson et al. 2020; Docherty et al. 2020; Simons et al. 2020; Grundy et al. 2020).

Although increased ACE2 in the conducting airways was suggested to increase susceptibility to SARS-CoV-2 infection (Cai et al, 2020); others have suggested elevated ACE2 in the distal airways or the pulmonary vasculature could be beneficial during the later stages of COVID-19 and mooted upregulation of ACE2 as a rational therapeutic goal (Chaudhry et al. 2020; Vaduganathan et al. 2020; Kuba et al. 2005; Monteil et al. 2020; Verdecchia et al. 2020).

We have now explored the link between smoking, ACE2 and SARS-CoV-2 infection *in vitro* using differentiated primary human bronchial airway epithelial cells (HBECs) at the air-liquid interface (ALI). Since these and similar nasal cells are the putative first sites of entry for SARS-CoV-2, we sought to expose these *in vitro* human “airways” to SARS-CoV-2 with and without cigarette smoke extract to understand the impact of smoking on the earliest stage of *in vivo* infection.

We demonstrate that cigarette smoke induces ACE2 expression in HBECs using multiple approaches – quantitative PCR, immunofluorescence and western blotting. This is consistent with the molecular epidemiology data linking ACE2 expression and smoking. Given the difficulties reproducing detection of ACE2 protein in clinical specimens (Aguiar et al. 2020; Hikmet et al. 2020; Ortiz et al. 2020), it is particularly important to have demonstrated expression of ACE2 protein and its localisation at the cell surface – where it has the capacity to act as a receptor for SARS-CoV-2. Importantly, in our experiments, increased expression of ACE2 in response to CSE does not significantly alter the percentage of cells infected. This is consistent with the epidemiological data suggesting smoking is not a major risk factor for infection.

One explanation for the lack of impact of CSE on HBEC infection despite elevation of ACE2 was that it upregulates the truncated isoform of ACE2 – dACE2 - lacking the SARS-CoV-2 binding domain. However, we demonstrate that CSE mainly induces expression of the full-length isoform, despite the cells retaining the potential to upregulate dACE2 in response to IFNα.

Our results differ from recently published data that did not detect an increase in ACE2 mRNA/protein in response to cigarette smoke but nevertheless suggested that smoking increases viral infection (Purkayastha et al. 2020). This discrepancy may reflect differences in the smoking exposure protocols used. In our experiments we added cigarette smoke extract to the basal chamber of the transwell while cells are maintained at ALI. This is a well-established technique and has been used extensively in respiratory and cardiovascular research (Schamberger et al. 2015; Ito, Ishimori, and Ishikawa 2018). Further, it recapitulated the increase in ACE2 reported in clinical specimens. Purkayastha *et al* took an alternative approach – exposure of the entire differentiated cell culture to smoke for a brief period of 3 minutes/day for 4 days. This protocol was modelled on a study by Gindele et al who adapted a customised, bespoke aerosol toxin exposure system to expose small airway epithelial cells to relatively higher cigarette smoke doses for longer duration and at repeated time-points during differentiation at ALI (Gindele et al. 2020). Their results demonstrated a clear induction of ACE2 (**Supplementary Figure 8**) also reported by Smith et al (Smith et al. 2020). In contrast, Purkayastha et al used a standard vacuum chamber with modest exposure doses and did not demonstrate ACE2 induction (Purkayastha et al. 2020). Of note, neither exposure model led to CSE-associated toxicity.

There has been a keen interest in modulating ACE2 expression to influence the course of COVID-19. This is a complex area, as noted by others (Michaud et al. 2020; Verdecchia et al. 2020; Zhang, Penninger, et al. 2020). One hypothesis is that reducing ACE2 expression in the upper airways may protect against SARS-CoV-2 infection. This is consistent with observations that transduction with ACE2 increases cellular susceptibility to infection albeit in models where the overexpression of ACE2 is at unrepresentative supraphysiological levels (McCray et al. 2007; Yang et al. 2007). It is critical to distinguish between the conducting and distal airways (alveolar units) when considering the impact of ACE2 expression. These cell types have different stem cell precursors, different functions and distinct responses to specific stimuli (Stripp and Shapiro 2006). Any potential enthusiasm for reducing ACE2 in the conducting airway where SARS-CoV-2 first infects cells must be tempered by a significant body of evidence suggesting higher ACE2 may have a protective impact in the distal airway and protect against acute lung injury/adult respiratory distress syndrome (Michaud et al. 2020; Verdecchia et al. 2020; Imai et al. 2005). Therefore, an ideal therapeutic in severe COVID-19 may reduce ACE2 in the conducting airways but increase its expression in the alveolar units. Although this is unlikely to be an achievable goal, our data demonstrating that an elevation of flACE2 has no impact on cellular infection is consistent with the notion that expression of flACE2 is not the key factor limiting SARS-CoV-2 infection of conducting airway cells.

In COPD, NRF2 agonists would be a theoretically attractive way to induce the key antioxidant response as well as modulate ACE2 expression. We show that oltipraz, an NRF2 agonist already in Phase 3 clinical trial for other indications, increased rather than decreased ACE2 in the HBECs. Nicotine, a component of cigarette smoke, is also being actively assessed in clinical trials as a potential protective agent in COVID-19 infection (ClinicalTrials.gov Identifier NCT04583410), (Farsalinos et al. 2020). This was based on early epidemiological observations suggesting smokers may be protected against SARS-CoV-2 infection as well as preclinical studies on submerged bronchial epithelial cells suggesting nicotine may increase ACE2 mRNA expression (Russo et al. 2020). It is uncertain how to translate observations on ACE2 mRNA in undifferentiated airway cells into a clear understanding of the receptor expression at the apical surface of airway epithelial cells. Our data show that nicotine does not significantly alter ACE2 or CHRNA7 mRNA expression in differentiated HBECs after 48 hours treatment.

Our studies are limited by the focus on the conducting airways and therefore address the initial phases of infection rather than the later stage at which individuals are admitted to hospital and which are important for COVID-19 morbidity and mortality. Further, smoking has important systemic impacts that cannot be modelled in ALI epithelial cultures. In that context it is possible that CSE in the basal media more closely mimics the sustained systemic effects of smoking. In terms of assaying cellular infection, differentiated airway ALI models have profound advantages but it is possible that subtle changes in susceptibility to infection are not detected because of the well-to-well variation typical of experiments using primary cells.

Overall, our data are entirely consistent with the documented epidemiology of SARS-CoV-2 infection. Individuals with chronic respiratory or cardiovascular disease associated are more vulnerable to severe COVID-19. However, current smokers have a similar susceptibility to SARS-CoV-2 infection as the general population. We show that the airway epithelial response to cigarette smoke is associated with an increase in full length ACE2 – the key SARS-CoV-2 receptor - but not an increased susceptibility to cellular infection. Therapeutic strategies that increase ACE2 receptor expression in the conducting airways are unlikely to increase cellular infection.

## Methods

### Primary human bronchial epithelial cell (HBEC) culture and other cell line culture

Primary human bronchial epithelial cells (HBECs) derived from a non-smoking donor (Cat# CC-2540, male; Lonza; Donor 1) or derived directly from a patient at Cambridge University Hospitals NHS Trust (Research Ethics Committee Reference 19/SW/0152; Donor 2) were expanded using PneumaCult™-Ex Plus Medium (Cat# 05040; Stemcell) supplemented with Penicillin (100 I.U./ml)-Streptomycin (100 μg/ml). All experiments were performed using cells at passage 3. All experiments in used cells expanded from Donor 1 except when stated otherwise.

Cell lines including A549 (ATCC; Cat# CCL-185, male), Calu3 (ATCC; Cat# HTB-55, male) HEK293T (ATCC; Cat# CRL-3216, female) (A549-ACE2, HEK293T-ACE2) have been used as negative or positive controls. HEK293T lines were maintained in RPMI supplemented with 10% FBS, 2 mM L-glutamine, pH 7.5, and 1 mM sodium pyruvate at 37°C in a 5% CO2. A549 and Calu-3 cell lines were cultured in Dulbecco’s Modified Eagle’s Medium (DMEM) and Eagle’s Minimal Essential Medium (EMEM) respectively, supplemented as specified above.

### Air-Liquid Interface (ALI) Culture

Briefly, 1 × 10^5^ of expanded primary HBECs at passage 3 in 200 l of supplemented PneumaCult™-Ex Plus Media were seeded in the apical chamber of a 24-well Transwell® insert with 0.4μM pore (Cat# 353095, Falcon) pre-coated with Rat tail Type I collagen (Cat# 354236, Corning) with 500 μl of PneumaCult™-Ex Plus Media in the basolateral chamber. The following day, both apical and basolateral chambers underwent a media change (200 μl and 500 μl, respectively). After two days of submerged culture, media from the apical chamber was removed to establish the air-liquid interface (ALI day 0) whilst media in the basolateral chamber was replaced with 500 μl HBEC ALI differentiation medium (PneumaCult™-ALI Medium, Cat# 05021; Stemcell). Basolateral media was changed every 2-3 days and apical surface washed with warm PBS twice a week to remove any build-up of mucous and secretions. Cultures were allowed to differentiate for at least 28 days before being used for any experiments.

### Cigarette Smoke Extract (CSE) Generation and Treatment

Cigarette smoke extract (CSE) was prepared, filter sterilised using 0.20 μm filter and used within 30 mins of generation. CSE was generated by smoking two Kentucky reference cigarettes and bubbling the generated smoke through 25 ml ALI media at a rate of 100 ml/min. Each cigarette took roughly 6 mins to burn. This solution is regarded as “100% CSE” and was diluted with ALI media to generate a 10% working solution. Cells were treated with 10% CSE for 48 hours before being treated with SARS-CoV-2, harvested or fixed for further analysis.

### SARS-CoV-2 infection

The clinical isolate of SARS-CoV-2 viruses used in this study were SARS-CoV-2/human/Liverpool/REMRQ0001/2020(Lineage B.29) (Chu et al. 2020) (Patterson et al. 2020) and SARS-CoV-2 England/ATACCC 174/2020 (Lineage B.1.1.7). Stocks were sequenced before use and the consensus matched the expected sequence exactly. Viral titre was determined by 50% tissue culture infectious dose (TCID50) in Huh7-ACE2 cells.

For infection, the indicated dose of virus was diluted in PBS to a final volume of 50 µL and added to the apical chamber of the transwell of differentiated HBEC-ALI cultures for 2-3 hours prior to removal. At 72 hours post-infection HBEC-ALI apical surfaces were washed once with PBS, dissociated with TrypLE, and fixed in 4% formaldehyde for 15 minutes. Fixed cells were washed and incubated for 15 minutes at room temperature in Perm/Wash buffer (BD #554723). Permeabilised cells were pelleted, stained for 15 minutes at room temperature in 100 µL of sheep anti-SARS-CoV-2 nucleocapsid antibody (MRC-PPU, DA114) at a concentration of 0.7 µg/mL, washed and incubated in 100 µL AF488 donkey anti-sheep (Jackson ImmunoResearch #713-545-147) at a concentration of 2 µg/mL for 15 minutes at room temperature. Stained cells were pelleted and fluorescence staining analysed on a BD Fortessa flow cytometer.

### Immunofluorescence

ALI cultures were washed three times with PBS and fixed using 4% paraformaldehyde (PFA) for 15 minutes at room temperature before permeabilization with 0.3% Triton-X for 15 minutes. Cells were blocked for 1 hour in 5% Normal goat serum/1% Bovine serum albumin (BSA) at room temperature. Primary antibodies; anti-ACE2 antibody was initially Abcam 228349 but was discontinued part-way through this study and was then replaced with 21115-1-AP (Proteintech); Acetylated tubulin (T7451; Sigma); Muc5AC (MA5-12178; Invitrogen), SARS-CoV / SARS-CoV-2 (COVID-19) spike antibody [1A9] (GTX632604; Genetex), SARS-CoV-2 (COVID-19) Nucleocapsid antibody DA114 (MRC PPU) were added and incubated at 4 degrees overnight. Following several washes with Phosphate Buffered Saline and Tween-20 (PBS-T), cultures were incubated with secondary antibodies for 1 hour in the dark at room temperature before being washed a further three times before Hoechst staining (100 μg/ml) and mounting. Confocal images were captured using a Nikon C2 Confocal Microscope, magnification ×40 oil. Composite images were generated and analysed using Fiji. Immunofluorescent images were also captured using a Cellomics Arrayscan (ThermoFisher Scientific VTI) using 64 fields of view/transwell at x10 magnification and analysed using HCS. Studio 2.0 Client Software. Results are expressed as percent of infected cells according to AF488 positive staining. For Figure 5E (Donor 1 only) the controls were shared with a published experiment ( )

### qRT-PCR

RNA was extracted using RNeasy Mini Kit (Qiagen) according to manufacturer’s instructions and quantified using a NanoDrop Spectrophotometer (ThermoFisher). cDNA synthesis was performed using a High-Capacity cDNA Reverse Transcription Kit (ThermoFisher). qRT-PCR was performed using Fast SYBR® Green Mix (ThermoFisher) alongside the following primers used for detecting expression of genes of interest: ACE2 Forward (5’-3’): CGAAGCCGAAGACCTGTTCTA, Reverse (5’-3’): GGGCAAGTGTGGACTGTTCC; dACE2 Forward (5’-3’): GGAAGCAGGCTGGGACAAA, Reverse(5’-3’): AGCTGTCAGGAAGTCGTCCATT; TBP (House keeper) Forward (5’-3’): AGTGAAGAACAGTCCAGACTG, Reverse (5’-3’): CCAGGAAATAACTCTGGCTCAT; TMPRSS2 Forward (5’-3’): CTGCTGGATTTCCGGGTG, Reverse (5’-3’) TTCTGAGGTCTTCCCTTTCTCCT; FOXJ1 Forward (5’-3’): TGGATCACGGACAACTTCTGCTA, Reverse (5’-3’) CACTTGTTCCAGAGACAGGTTGTGG; MUC5B Forward (5’-3’): CCTGAAGTCTTCCCCAGCAG, Reverse (5’-3’) GCATAGAATTGGCAGCCAGC. Samples were run in technical triplicates on a StepOne machine and relative differences in expression were determined using the comparative ⊗C_T_ method and TBP used as the endogenous house-keeping control.

### Western Blotting

Recombinant Anti-ACE2 antibodies (Cat# ab108209; Abcam: N-terminal) and (Cat# ab15348; Abcam: C-terminal), alpha-tubulin (Cat# sc-32293; Santa-Cruz) and beta-actin (Cat# sc-69879; Santa-Cruz) were used for ACE2/dACE2, alpha-tubulin detection and beta-actin, respectively.

### Apoptosis detection

ALI cultures exposed to CSE or control media were washed three times with PBS and detached from the transwell membrane with accutase. Apoptotic cells was detected by concurrent staining with annexin V–APC and PI (Cat# 88-8007-72, eBioscience) and their far-red and red fluorescence was measured by flow cytometry (Fortessa LSR, BD).

### Quantification and Statistical Analysis

Statistical analyses of mRNA expression assays and infection quantification data were performed using Prism 8 software (GraphPad Software). P values were calculated using a two-tailed, Mann Whitney *U*-test unless stated otherwise. P values were noted as follows: ns, not significant; *, P < 0.05; **, P < 0.01; ***, P < 0.001. Error bars represent the mean +/-standard error of the mean unless stated otherwise.

## Acknowledgements

SARS-CoV-2/human/Liverpool/REMRQ0001/2020 was a kind gift from Lance Turtle (University of Liverpool) and David Matthews and Andrew Davidson (University of Bristol). SARS-CoV-2 England/ATACCC 174/2020 was a kind gift from Greg Towers (University College London), and we are also grateful to Ajit Lalvani, Jake Dunning, Maria Zambon and colleagues at Public Health England and Giada Mattiuzzo at the National Institute for Biological Standards and Controls and Wendy Barclay and Jonathan Brown and all colleagues in the United Kingdom Research Institute funded collaboration Genotype to Phenotype. Sheep anti-SARS-CoV-2 nucleoprotein antibody (DA114) was a kind gift from Paul Davies (obtained from MRC PPU Reagents and Services, University of Dundee). We gratefully acknowledge the support from Ms Jacqui Galloway in establishing the primary cells from patients. We also gratefully acknowledge advice and discussions from Dr Stephen White, Department of Life Sciences, Manchester Metropolitan University regarding generation and use of cigarette smoke extract.

We are grateful for the generous support of the UKRI COVID Immunology Consortium, Addenbrooke’s Charitable Trust (15/20A) and the NIHR Cambridge Biomedical Research Centre. This work was supported by a Wellcome Trust Principal Research Fellowship (084957/Z/08/Z) and MRC research grant MR/V011561/1 to P.J.L. This work was supported by the NC3Rs NC/S001204/1 project grant and the Roy Castle Lung Cancer Foundation grant (2015/10/McCaughan) to FM.

This paper presents independent research supported by the NIHR Cambridge BRC. The NIHR Cambridge Biomedical Research Centre (BRC) is a partnership between Cambridge University Hospitals NHS Foundation Trust and the University of Cambridge, funded by the National Institute for Health Research (NIHR). The views expressed are those of the author(s) and not necessarily those of the NIHR or the Department of Health and Social Care.

The authors declare no competing interests.

## REFERENCES

Aguiar, J. A., B. J. Tremblay, M. J. Mansfield, O. Woody, B. Lobb, A. Banerjee, A. Chandiramohan, N. Tiessen, Q. Cao, A. Dvorkin-Gheva, S. Revill, M. S. Miller, C. Carlsten, L. Organ, C. Joseph, A. John, P. Hanson, R. C. Austin, B. M. McManus, G. Jenkins, K. Mossman, K. Ask, A. C. Doxey, and J. A. Hirota. 2020. ‘Gene expression and in situ protein profiling of candidate SARS-CoV-2 receptors in human airway epithelial cells and lung tissue’, Eur Respir J, 56.

Aliee, H., F. Massip, C. Qi, M. Stella de Biase, J. L. van Nijnatten, E. T. G. Kersten, N. Z. Kermani, B. Khuder, J. M. Vonk, R. C. H. Vermeulen, M. Neighbors, G. W. Tew, M. Grimbaldeston, N. H. T. Ten Hacken, S. Hu, Y. Guo, X. Zhang, K. Sun, P. S. Hiemstra, B. A. Ponder, M. J. Makela, K. Malmstrom, R. C. Rintoul, P. A. Reyfman, F. J. Theis, C. A. Brandsma, I. Adcock, W. Timens, C. J. Xu, M. van den Berge, R. F. Schwarz, G. H. Koppelman, M. C. Nawijn, and A. Faiz. 2020. ‘Determinants of SARS-CoV-2 receptor gene expression in upper and lower airways’, medRxiv.

Alqahtani, J. S., T. Oyelade, A. M. Aldhahir, S. M. Alghamdi, M. Almehmadi, A. S. Alqahtani, S. Quaderi, S. Mandal, and J. R. Hurst. 2020. ‘Prevalence, Severity and Mortality associated with COPD and Smoking in patients with COVID-19: A Rapid Systematic Review and Meta-Analysis’, PLoS One, 15: e0233147.

Bindom, S. M., C. P. Hans, H. Xia, A. H. Boulares, and E. Lazartigues. 2010. ‘Angiotensin I-converting enzyme type 2 (ACE2) gene therapy improves glycemic control in diabetic mice’, Diabetes, 59: 2540–8.

Blanco-Melo, D., B. E. Nilsson-Payant, W. C. Liu, S. Uhl, D. Hoagland, R. Moller, T. X. Jordan, K. Oishi, M. Panis, D. Sachs, T. T. Wang, R. E. Schwartz, J. K. Lim, R. A. Albrecht, and B. R. tenOever. 2020. ‘Imbalanced Host Response to SARS-CoV-2 Drives Development of COVID-19’, Cell, 181: 1036–45 e9.

Blume, C., C. L. Jackson, C. M. Spalluto, J. Legebeke, L. Nazlamova, F. Conforti, J. M. Perotin, M. Frank, J. Butler, M. Crispin, J. Coles, J. Thompson, R. A. Ridley, L. S. N. Dean, M. Loxham, S. Reikine, A. Azim, K. Tariq, D. A. Johnston, P. J. Skipp, R. Djukanovic, D. Baralle, C. J. McCormick, D. E. Davies, J. S. Lucas, G. Wheway, and V. Mennella. 2021. ‘A novel ACE2 isoform is expressed in human respiratory epithelia and is upregulated in response to interferons and RNA respiratory virus infection’, Nat Genet, 53: 205–14.

Brake, S. J., K. Barnsley, W. Lu, K. D. McAlinden, M. S. Eapen, and S. S. Sohal. 2020. ‘Smoking Upregulates Angiotensin-Converting Enzyme-2 Receptor: A Potential Adhesion Site for Novel Coronavirus SARS-CoV-2 (Covid-19)’, J Clin Med, 9.

Cai, G., Y. Bosse, F. Xiao, F. Kheradmand, and C. I. Amos. 2020. ‘Tobacco Smoking Increases the Lung Gene Expression of ACE2, the Receptor of SARS-CoV-2’, Am J Respir Crit Care Med, 201: 1557–59.

Chaudhry, F., S. Lavandero, X. Xie, B. Sabharwal, Y. Y. Zheng, A. Correa, J. Narula, and P. Levy. 2020. ‘Manipulation of ACE2 expression in COVID-19’, Open Heart, 7.

Chu, H., J. F. Chan, T. T. Yuen, H. Shuai, S. Yuan, Y. Wang, B. Hu, C. C. Yip, J. O. Tsang, X. Huang, Y. Chai, D. Yang, Y. Hou, K. K. Chik, X. Zhang, A. Y. Fung, H. W. Tsoi, J. P. Cai, W. M. Chan, J. D. Ip, A. W. Chu, J. Zhou, D. C. Lung, K. H. Kok, K.K. To, O. T. Tsang, K. H. Chan, and K. Y. Yuen. 2020. ‘Comparative tropism, replication kinetics, and cell damage profiling of SARS-CoV-2 and SARS-CoV with implications for clinical manifestations, transmissibility, and laboratory studies of COVID-19: an observational study’, Lancet Microbe, 1: e14–e23.

Chung, M. K., S. Karnik, J. Saef, C. Bergmann, J. Barnard, M. M. Lederman, J. Tilton, F. Cheng, C. V. Harding, J. B. Young, N. Mehta, S. J. Cameron, K. R. McCrae, A. H. Schmaier, J. D. Smith, A. Kalra, S. K. Gebreselassie, G. Thomas, E. S. Hawkins, and L. G. Svensson. 2020. ‘SARS-CoV-2 and ACE2: The biology and clinical data settling the ARB and ACEI controversy’, EBioMedicine, 58: 102907.

Daly, J. L., B. Simonetti, K. Klein, K. E. Chen, M. K. Williamson, C. Anton-Plagaro, D. K. Shoemark, L. Simon-Gracia, M. Bauer, R. Hollandi, U. F. Greber, P. Horvath, R. B. Sessions, A. Helenius, J. A. Hiscox, T. Teesalu, D. A. Matthews, A. D. Davidson, B. M. Collins, P. J. Cullen, and Y. Yamauchi. 2020. ‘Neuropilin-1 is a host factor for SARS-CoV-2 infection’, Science, 370: 861–65.

Docherty, A. B., E. M. Harrison, C. A. Green, H. E. Hardwick, R. Pius, L. Norman, K. A. Holden, J. M. Read, F. Dondelinger, G. Carson, L. Merson, J. Lee, D. Plotkin, L. Sigfrid, S. Halpin, C. Jackson, C. Gamble, P. W. Horby, J. S. Nguyen-Van-Tam, A. Ho, C. D. Russell, J. Dunning, P. J. Openshaw, J. K. Baillie, M. G. Semple, and Isaric C. investigators. 2020. ‘Features of 20 133 UK patients in hospital with covid-19 using the ISARIC WHO Clinical Characterisation Protocol: prospective observational cohort study’, BMJ, 369: m1985.

Duffney, P. F., C. E. McCarthy, A. Nogales, T. H. Thatcher, L. Martinez-Sobrido, R. P. Phipps, and P. J. Sime. 2018. ‘Cigarette smoke dampens antiviral signaling in small airway epithelial cells by disrupting TLR3 cleavage’, Am J Physiol Lung Cell Mol Physiol, 314: L505–L13.

Eddleston, J., R. U. Lee, A. M. Doerner, J. Herschbach, and B. L. Zuraw. 2011. ‘Cigarette smoke decreases innate responses of epithelial cells to rhinovirus infection’, Am J Respir Cell Mol Biol, 44: 118–26.

Farsalinos, K., A. Angelopoulou, N. Alexandris, and K. Poulas. 2020. ‘COVID-19 and the nicotinic cholinergic system’, Eur Respir J, 56.

Gebel, S., S. Diehl, J. Pype, B. Friedrichs, H. Weiler, J. Schuller, H. Xu, K. Taguchi, M. Yamamoto, and T. Muller. 2010. ‘The transcriptome of Nrf2-/-mice provides evidence for impaired cell cycle progression in the development of cigarette smoke-induced emphysematous changes’, Toxicol Sci, 115: 238–52.

Gindele, J. A., T. Kiechle, K. Benediktus, G. Birk, M. Brendel, F. Heinemann, C. T. Wohnhaas, M. LeBlanc, H. Zhang, Y. Strulovici-Barel, R. G. Crystal, M. J. Thomas, B. Stierstorfer, K. Quast, and J. Schymeinsky. 2020. ‘Intermittent exposure to whole cigarette smoke alters the differentiation of primary small airway epithelial cells in the air-liquid interface culture’, Sci Rep, 10: 6257.

Grundy, E. J., T. Suddek, F. T. Filippidis, A. Majeed, and S. Coronini-Cronberg. 2020. ‘Smoking, SARS-CoV-2 and COVID-19: A review of reviews considering implications for public health policy and practice’, Tob Induc Dis, 18: 58.

Guo, F. R. 2020. ‘Smoking links to the severity of COVID-19: An update of a meta-analysis’, J Med Virol, 92: 2304–05.

Hamming, I., W. Timens, M. L. Bulthuis, A. T. Lely, G. Navis, and H. van Goor. 2004. ‘Tissue distribution of ACE2 protein, the functional receptor for SARS coronavirus. A first step in understanding SARS pathogenesis’, J Pathol, 203: 631–7.

Harmer, D., M. Gilbert, R. Borman, and K. L. Clark. 2002. ‘Quantitative mRNA expression profiling of ACE 2, a novel homologue of angiotensin converting enzyme’, FEBS Lett, 532: 107–10.

Hikmet, F., L. Mear, A. Edvinsson, P. Micke, M. Uhlen, and C. Lindskog. 2020. ‘The protein expression profile of ACE2 in human tissues’, Mol Syst Biol, 16: e9610.

Hoffmann, M., H. Kleine-Weber, S. Schroeder, N. Kruger, T. Herrler, S. Erichsen, T. S. Schiergens, G. Herrler, N. H. Wu, A. Nitsche, M. A. Muller, C. Drosten, and S. Pohlmann. 2020. ‘SARS-CoV-2 Cell Entry Depends on ACE2 and TMPRSS2 and Is Blocked by a Clinically Proven Protease Inhibitor’, Cell, 181: 271–80 e8.

Hopkinson, N. S., N. Rossi, J. El-Sayed Moustafa, A. A. Laverty, J. K. Quint, M. Freidin, A. Visconti, B. Murray, M. Modat, S. Ourselin, K. Small, R. Davies, J. Wolf, T. D. Spector, C. J. Steves, and M. Falchi. 2021. ‘Current smoking and COVID-19 risk: results from a population symptom app in over 2.4 million people’, Thorax.

Hou, Y. J., K. Okuda, C. E. Edwards, D. R. Martinez, T. Asakura, K. H. Dinnon, 3rd, T. Kato, R. E. Lee, B. L. Yount, T. M. Mascenik, G. Chen, K. N. Olivier, A. Ghio, L. V. Tse, S. R. Leist, L. E. Gralinski, A. Schafer, H. Dang, R. Gilmore, S. Nakano, L. Sun, M. L. Fulcher, A. Livraghi-Butrico, N. I. Nicely, M. Cameron, C. Cameron, D. J. Kelvin, A. de Silva, D. M. Margolis, A. Markmann, L. Bartelt, R. Zumwalt, F. J. Martinez, S. P. Salvatore, A. Borczuk, P. R. Tata, V. Sontake, A. Kimple, I. Jaspers, W.K. O’Neal, S. H. Randell, R. C. Boucher, and R. S. Baric. 2020. ‘SARS-CoV-2 Reverse Genetics Reveals a Variable Infection Gradient in the Respiratory Tract’, Cell, 182: 429–46 e14.

Hung, Y. H., W. Y. Hsieh, J. S. Hsieh, F. C. Liu, C. H. Tsai, L. C. Lu, C. Y. Huang, C. L. Wu, and C. S. Lin. 2016. ‘Alternative Roles of STAT3 and MAPK Signaling Pathways in the MMPs Activation and Progression of Lung Injury Induced by Cigarette Smoke Exposure in ACE2 Knockout Mice’, Int J Biol Sci, 12: 454–65.

Imai, Y., K. Kuba, S. Rao, Y. Huan, F. Guo, B. Guan, P. Yang, R. Sarao, T. Wada, H. Leong-Poi, M. A. Crackower, A. Fukamizu, C. C. Hui, L. Hein, S. Uhlig, A. S. Slutsky, C. Jiang, and J. M. Penninger. 2005. ‘Angiotensin-converting enzyme 2 protects from severe acute lung failure’, Nature, 436: 112–6.

Ito, S., K. Ishimori, and S. Ishikawa. 2018. ‘Effects of repeated cigarette smoke extract exposure over one month on human bronchial epithelial organotypic culture’, Toxicol Rep, 5: 864–70.

Jacobs, M., H. P. Van Eeckhoutte, S. R. A. Wijnant, W. Janssens, G. F. Joos, G. G. Brusselle, and K. R. Bracke. 2020. ‘Increased expression of ACE2, the SARS-CoV-2 entry receptor, in alveolar and bronchial epithelium of smokers and COPD subjects’, Eur Respir J, 56.

Jia, H. P., D. C. Look, L. Shi, M. Hickey, L. Pewe, J. Netland, M. Farzan, C. Wohlford-Lenane, S. Perlman, and P. B. McCray, Jr. 2005. ‘ACE2 receptor expression and severe acute respiratory syndrome coronavirus infection depend on differentiation of human airway epithelia’, J Virol, 79: 14614–21.

Jiang, F., J. Yang, Y. Zhang, M. Dong, S. Wang, Q. Zhang, F. F. Liu, K. Zhang, and C. Zhang. 2014. ‘Angiotensin-converting enzyme 2 and angiotensin 1-7: novel therapeutic targets’, Nat Rev Cardiol, 11: 413–26.

Kuba, K., Y. Imai, S. Rao, H. Gao, F. Guo, B. Guan, Y. Huan, P. Yang, Y. Zhang, W. Deng, L. Bao, B. Zhang, G. Liu, Z. Wang, M. Chappell, Y. Liu, D. Zheng, A. Leibbrandt, T. Wada, A. S. Slutsky, D. Liu, C. Qin, C. Jiang, and J. M. Penninger. 2005. ‘A crucial role of angiotensin converting enzyme 2 (ACE2) in SARS coronavirus-induced lung injury’, Nat Med, 11: 875–9.

Lan, J., J. Ge, J. Yu, S. Shan, H. Zhou, S. Fan, Q. Zhang, X. Shi, Q. Wang, L. Zhang, and X. Wang. 2020. ‘Structure of the SARS-CoV-2 spike receptor-binding domain bound to the ACE2 receptor’, Nature, 581: 215–20.

Lee, I. T., T. Nakayama, C. T. Wu, Y. Goltsev, S. Jiang, P. A. Gall, C. K. Liao, L. C. Shih, C. M. Schurch, D. R. McIlwain, P. Chu, N. A. Borchard, D. Zarabanda, S. S. Dholakia, A. Yang, D. Kim, H. Chen, T. Kanie, C. D. Lin, M. H. Tsai, K. M. Phillips, R. Kim, J. B. Overdevest, M. A. Tyler, C. H. Yan, C. F. Lin, Y. T. Lin, D. T. Bau, G. J. Tsay, Z. M. Patel, Y. A. Tsou, A. Tzankov, M. S. Matter, C. J. Tai, T. H. Yeh, P. H. Hwang, G. P. Nolan, J. V. Nayak, and P. K. Jackson. 2020. ‘ACE2 localizes to the respiratory cilia and is not increased by ACE inhibitors or ARBs’, Nat Commun, 11: 5453.

Leung, J. M., C. X. Yang, A. Tam, T. Shaipanich, T. L. Hackett, G. K. Singhera, D. R. Dorscheid, and D. D. Sin. 2020. ‘ACE-2 expression in the small airway epithelia of smokers and COPD patients: implications for COVID-19’, Eur Respir J, 55.

Li, F., W. Li, M. Farzan, and S. C. Harrison. 2005. ‘Structure of SARS coronavirus spike receptor-binding domain complexed with receptor’, Science, 309: 1864–8.

Li, W., M. J. Moore, N. Vasilieva, J. Sui, S. K. Wong, M. A. Berne, M. Somasundaran, J. L. Sullivan, K. Luzuriaga, T. C. Greenough, H. Choe, and M. Farzan. 2003. ‘Angiotensin-converting enzyme 2 is a functional receptor for the SARS coronavirus’, Nature, 426: 450–4.

Lukassen, S., R. L. Chua, T. Trefzer, N. C. Kahn, M. A. Schneider, T. Muley, H. Winter, M. Meister, C. Veith, A. W. Boots, B. P. Hennig, M. Kreuter, C. Conrad, and R. Eils. 2020. ‘SARS-CoV-2 receptor ACE2 and TMPRSS2 are primarily expressed in bronchial transient secretory cells’, EMBO J, 39: e105114.

McCray, P. B., Jr., L. Pewe, C. Wohlford-Lenane, M. Hickey, L. Manzel, L. Shi, J. Netland, H. P. Jia, C. Halabi, C. D. Sigmund, D. K. Meyerholz, P. Kirby, D. C. Look, and S. Perlman. 2007. ‘Lethal infection of K18-hACE2 mice infected with severe acute respiratory syndrome coronavirus’, J Virol, 81: 813–21.

Michaud, V., M. Deodhar, M. Arwood, S. B. Al Rihani, P. Dow, and J. Turgeon. 2020. ‘ACE2 as a Therapeutic Target for COVID-19; its Role in Infectious Processes and Regulation by Modulators of the RAAS System’, J Clin Med, 9.

Monteil, V., H. Kwon, P. Prado, A. Hagelkruys, R. A. Wimmer, M. Stahl, A. Leopoldi, E. Garreta, C. Hurtado Del Pozo, F. Prosper, J. P. Romero, G. Wirnsberger, H. Zhang, A. S. Slutsky, R. Conder, N. Montserrat, A. Mirazimi, and J. M. Penninger. 2020. ‘Inhibition of SARS-CoV-2 Infections in Engineered Human Tissues Using Clinical-Grade Soluble Human ACE2’, Cell, 181: 905–13 e7.

Ng, K. W., J. Attig, W. Bolland, G. R. Young, J. Major, A. G. Wrobel, S. Gamblin, A. Wack, and G. Kassiotis. 2020. ‘Tissue-specific and interferon-inducible expression of nonfunctional ACE2 through endogenous retroelement co-option’, Nat Genet, 52: 1294–302.

Niu, M. J., J. K. Yang, S. S. Lin, X. J. Ji, and L. M. Guo. 2008. ‘Loss of angiotensin-converting enzyme 2 leads to impaired glucose homeostasis in mice’, Endocrine, 34: 56–61.

Olagnier, D., E. Farahani, J. Thyrsted, J. Blay-Cadanet, A. Herengt, M. Idorn, A. Hait, B. Hernaez, A. Knudsen, M. B. Iversen, M. Schilling, S. E. Jorgensen, M. Thomsen, L. S. Reinert, M. Lappe, H. D. Hoang, V. H. Gilchrist, A. L. Hansen, R. Ottosen, C. G. Nielsen, C. Moller, D. van der Horst, S. Peri, S. Balachandran, J. Huang, M. Jakobsen, E. B. Svenningsen, T. B. Poulsen, L. Bartsch, A. L. Thielke, Y. Luo, T. Alain, J. Rehwinkel, A. Alcami, J. Hiscott, T. H. Mogensen, S. R. Paludan, and C. K. Holm. 2020. ‘SARS-CoV2-mediated suppression of NRF2-signaling reveals potent antiviral and anti-inflammatory activity of 4-octyl-itaconate and dimethyl fumarate’, Nat Commun, 11: 4938.

Onabajo, O. O., A. R. Banday, W. Yan, A. Obajemu, M. L. Stanifer, D. M. Santer, O. Florez-Vargas, H. Piontkivska, J. Vargas, C. Kee, D. L. J. Tyrrell, J. L. Mendoza, S. Boulant, and L. Prokunina-Olsson. 2020. ‘Interferons and viruses induce a novel primate-specific isoform dACE2 and not the SARS-CoV-2 receptor ACE2’, bioRxiv.

Ortiz, M. E., A. Thurman, A. A. Pezzulo, M. R. Leidinger, J. A. Klesney-Tait, P. H. Karp, P. Tan, C. Wohlford-Lenane, P. B. McCray, Jr., and D. K. Meyerholz. 2020. ‘Heterogeneous expression of the SARS-Coronavirus-2 receptor ACE2 in the human respiratory tract’, EBioMedicine, 60: 102976.

Patanavanich, R., and S. A. Glantz. 2020. ‘Smoking Is Associated With COVID-19 Progression: A Meta-analysis’, Nicotine Tob Res, 22: 1653–56.

Patterson, E. I., T. Prince, E. R. Anderson, A. Casas-Sanchez, S. L. Smith, C. Cansado-Utrilla, L. Turtle, and G. L. Hughes. 2020. ‘Methods of inactivation of SARS-CoV-2 for downstream biological assays’, bioRxiv.

Purkayastha, A., C. Sen, G. Garcia, Jr., J. Langerman, D. W. Shia, L. K. Meneses, P. Vijayaraj, A. Durra, C. R. Koloff, D. R. Freund, J. Chi, T. M. Rickabaugh, A. Mulay, B. Konda, M. S. Sim, B. R. Stripp, K. Plath, V. Arumugaswami, and B. N. Gomperts. 2020. ‘Direct Exposure to SARS-CoV-2 and Cigarette Smoke Increases Infection Severity and Alters the Stem Cell-Derived Airway Repair Response’, Cell Stem Cell, 27: 869–75 e4.

Ravindra, N. G., M. M. Alfajaro, V. Gasque, N. C. Huston, H. Wan, K. Szigeti-Buck, Y. Yasumoto, A. M. Greaney, V. Habet, R. D. Chow, J. S. Chen, J. Wei, R. B. Filler, B. Wang, G. Wang, L. E. Niklason, R. R. Montgomery, S. C. Eisenbarth, S. Chen, A. Williams, A. Iwasaki, T. L. Horvath, E. F. Foxman, R. W. Pierce, A. M. Pyle, D. van Dijk, and C. B. Wilen. 2021. ‘Single-cell longitudinal analysis of SARS-CoV-2 infection in human airway epithelium identifies target cells, alterations in gene expression, and cell state changes’, PLoS Biol, 19: e3001143.

Ren, X., J. Glende, M. Al-Falah, V. de Vries, C. Schwegmann-Wessels, X. Qu, L. Tan, T. Tschernig, H. Deng, H. Y. Naim, and G. Herrler. 2006. ‘Analysis of ACE2 in polarized epithelial cells: surface expression and function as receptor for severe acute respiratory syndrome-associated coronavirus’, J Gen Virol, 87: 1691–95.

Rossato, M., L. Russo, S. Mazzocut, A. Di Vincenzo, P. Fioretto, and R. Vettor. 2020. ‘Current smoking is not associated with COVID-19’, Eur Respir J, 55.

Russo, P., S. Bonassi, R. Giacconi, M. Malavolta, C. Tomino, and F. Maggi. 2020. ‘COVID-19 and smoking: is nicotine the hidden link?’, Eur Respir J, 55.

Sachs, L. A., W. E. Finkbeiner, and J. H. Widdicombe. 2003. ‘Effects of media on differentiation of cultured human tracheal epithelium’, In Vitro Cell Dev Biol Anim, 39: 56–62.

Schaefer, I. M., R. F. Padera, I. H. Solomon, S. Kanjilal, M. M. Hammer, J. L. Hornick, and L. M. Sholl. 2020. ‘In situ detection of SARS-CoV-2 in lungs and airways of patients with COVID-19’, Mod Pathol, 33: 2104–14.

Schamberger, A. C., C. A. Staab-Weijnitz, N. Mise-Racek, and O. Eickelberg. 2015. ‘Cigarette smoke alters primary human bronchial epithelial cell differentiation at the air-liquid interface’, Sci Rep, 5: 8163.

Shajahan, A., S. Archer-Hartmann, N. T. Supekar, A. S. Gleinich, C. Heiss, and P. Azadi. 2020. ‘Comprehensive characterization of N- and O-glycosylation of SARS-CoV-2 human receptor angiotensin converting enzyme 2’, Glycobiology.

Shang, J., Y. Wan, C. Luo, G. Ye, Q. Geng, A. Auerbach, and F. Li. 2020. ‘Cell entry mechanisms of SARS-CoV-2’, Proc Natl Acad Sci U S A, 117: 11727–34.

Simons, D., L. Shahab, J. Brown, and O. Perski. 2020. ‘The association of smoking status with SARS-CoV-2 infection, hospitalization and mortality from COVID-19: a living rapid evidence review with Bayesian meta-analyses (version 7)’, Addiction.

Sims, A. C., R. S. Baric, B. Yount, S. E. Burkett, P. L. Collins, and R. J. Pickles. 2005. ‘Severe acute respiratory syndrome coronavirus infection of human ciliated airway epithelia: role of ciliated cells in viral spread in the conducting airways of the lungs’, J Virol, 79: 15511–24.

Smith, J. C., E. L. Sausville, V. Girish, M. L. Yuan, A. Vasudevan, K. M. John, and J. M. Sheltzer. 2020. ‘Cigarette Smoke Exposure and Inflammatory Signaling Increase the Expression of the SARS-CoV-2 Receptor ACE2 in the Respiratory Tract’, Dev Cell, 53: 514–29 e3.

Stripp, B. R., and S. D. Shapiro. 2006. ‘Stem cells in lung disease, repair, and the potential for therapeutic interventions: State-of-the-art and future challenges’, Am J Respir Cell Mol Biol, 34: 517–18.

Sungnak, W., N. Huang, C. Becavin, M. Berg, R. Queen, M. Litvinukova, C. Talavera-Lopez, H. Maatz, D. Reichart, F. Sampaziotis, K. B. Worlock, M. Yoshida, J. L. Barnes, and H. C. A. Lung Biological Network. 2020. ‘SARS-CoV-2 entry factors are highly expressed in nasal epithelial cells together with innate immune genes’, Nat Med, 26: 681–87.

Vaduganathan, M., O. Vardeny, T. Michel, J. J. V. McMurray, M. A. Pfeffer, and S. D. Solomon. 2020. ‘Renin-Angiotensin-Aldosterone System Inhibitors in Patients with Covid-19’, N Engl J Med, 382: 1653–59.

Verdecchia, P., C. Cavallini, A. Spanevello, and F. Angeli. 2020. ‘The pivotal link between ACE2 deficiency and SARS-CoV-2 infection’, Eur J Intern Med, 76: 14–20.

Walls, A. C., Y. J. Park, M. A. Tortorici, A. Wall, A. T. McGuire, and D. Veesler. 2020. ‘Structure, Function, and Antigenicity of the SARS-CoV-2 Spike Glycoprotein’, Cell, 181: 281–92 e6.

Wang, Q., Y. Zhang, L. Wu, S. Niu, C. Song, Z. Zhang, G. Lu, C. Qiao, Y. Hu, K. Y. Yuen, Q. Wang, H. Zhou, J. Yan, and J. Qi. 2020. ‘Structural and Functional Basis of SARS-CoV-2 Entry by Using Human ACE2’, Cell, 181: 894–904 e9.

Williamson, E. J., A. J. Walker, K. Bhaskaran, S. Bacon, C. Bates, C. E. Morton, H. J. Curtis, A. Mehrkar, D. Evans, P. Inglesby, J. Cockburn, H. I. McDonald, B. MacKenna, L. Tomlinson, I. J. Douglas, C. T. Rentsch, R. Mathur, A. Y. S. Wong, R. Grieve, D. Harrison, H. Forbes, A. Schultze, R. Croker, J. Parry, F. Hester, S. Harper, R. Perera, S. J. W. Evans, L. Smeeth, and B. Goldacre. 2020. ‘Factors associated with COVID-19-related death using OpenSAFELY’, Nature, 584: 430–36.

Yang, X. H., W. Deng, Z. Tong, Y. X. Liu, L. F. Zhang, H. Zhu, H. Gao, L. Huang, Y. L. Liu, C. M. Ma, Y. F. Xu, M. X. Ding, H. K. Deng, and C. Qin. 2007. ‘Mice transgenic for human angiotensin-converting enzyme 2 provide a model for SARS coronavirus infection’, Comp Med, 57: 450–9.

Yilin, Z., N. Yandong, and J. Faguang. 2015. ‘Role of angiotensin-converting enzyme (ACE) and ACE2 in a rat model of smoke inhalation induced acute respiratory distress syndrome’, Burns, 41: 1468–77.

Zamorano Cuervo, N., and N. Grandvaux. 2020. ‘ACE2: Evidence of role as entry receptor for SARS-CoV-2 and implications in comorbidities’, Elife, 9.

Zhang, H., J. M. Penninger, Y. Li, N. Zhong, and A. S. Slutsky. 2020. ‘Angiotensin-converting enzyme 2 (ACE2) as a SARS-CoV-2 receptor: molecular mechanisms and potential therapeutic target’, Intensive Care Med, 46: 586–90.

Zhang, Q., Y. Yue, H. Tan, Y. Liu, Y. Zeng, and L. Xiao. 2020. ‘Single Cell RNA-seq Data Analysis Reveals the Potential Risk of SARS-CoV-2 Infection Among Different Respiratory System Conditions’, Front Genet, 11: 942.

Zhao, Q., M. Meng, R. Kumar, Y. Wu, J. Huang, N. Lian, Y. Deng, and S. Lin. 2020. ‘The impact of COPD and smoking history on the severity of COVID-19: A systemic review and meta-analysis’, J Med Virol, 92: 1915–21.

Ziegler, C. G. K., S. J. Allon, S. K. Nyquist, I. M. Mbano, V. N. Miao, C. N. Tzouanas, Y. Cao, A. S. Yousif, J. Bals, B. M. Hauser, J. Feldman, C. Muus, M. H. Wadsworth, 2nd, S. W. Kazer, T. K. Hughes, B. Doran, G. J. Gatter, M. Vukovic, F. Taliaferro, B.E. Mead, Z. Guo, J. P. Wang, D. Gras, M. Plaisant, M. Ansari, I. Angelidis, H. Adler, J. M. S. Sucre, C. J. Taylor, B. Lin, A. Waghray, V. Mitsialis, D. F. Dwyer, K. M. Buchheit, J. A. Boyce, N. A. Barrett, T. M. Laidlaw, S. L. Carroll, L. Colonna, V. Tkachev, C. W. Peterson, A. Yu, H. B. Zheng, H. P. Gideon, C. G. Winchell, P. L. Lin, C. D. Bingle, S. B. Snapper, J. A. Kropski, F. J. Theis, H. B. Schiller, L. E. Zaragosi, P. Barbry, A. Leslie, H. P. Kiem, J. L. Flynn, S. M. Fortune, B. Berger, R.W. Finberg, L. S. Kean, M. Garber, A. G. Schmidt, D. Lingwood, A. K. Shalek, J. Ordovas-Montanes H. C. A. Lung Biological Network. Electronic address, and H. C. A. Lung Biological Network. 2020. ‘SARS-CoV-2 Receptor ACE2 Is an Interferon-Stimulated Gene in Human Airway Epithelial Cells and Is Detected in Specific Cell Subsets across Tissues’, Cell, 181: 1016–l35 e19.

Zou, X., K. Chen, J. Zou, P. Han, J. Hao, and Z. Han. 2020. ‘Single-cell RNA-seq data analysis on the receptor ACE2 expression reveals the potential risk of different human organs vulnerable to 2019-nCoV infection’, Front Med, 14: 185–92.

