## Supplementary Figures for "Cigarette smoke preferentially induces full length ACE2 exposure in primary human airway cells but does not alter susceptibility to SARS-CoV-2 infection"

**Suppl Figure 1 CSE upregulates *ACE2* expression relative to control wells.**

**A.** Differentiated HBECs (Donor 1) cultured at the ALI for 28 days were stained with antibodies against ACE2 (red, 21115-AP) and acetylated tubulin (green). **B.** Red fluorescence in the untreated well was reduced until ACE2 signal could not be detected and this threshold was then applied to CSE exposed wells. CSE exposed wells showed a persistent apical ACE2 signal. Data is representative of two independent experiments.

**Suppl Figure 2. Exposure to CSE induces ACE2 expression in differentiated HBECs derived from an individual with COPD (Donor 2)**

**A**. CSE exposure (10%) for 48 h increases *ACE2* expression in differentiated COPD-derived HBECs relative to untreated controls. RT-PCR data shows log2 relative fold-change in expression from n=4 independent experiments, (Mann-Whitney, *p<0.05). **B.** CSE exposure (10%) for 48 h also increases ACE2 protein expression relative to untreated control (Antibody: Ab15348). Calu-3 and A549-ACE2 are presented as the positive controls. Representative western blot from 4 independent experiments.

**Suppl Figure 3. Representative flow cytometric plots and histograms from CSE exposed and untreated HBEC cultures (Donor 1).** CSE exposure does not induce apoptosis in differentiated HBECs at ALI relative to control samples. HBECs at ALI exposed to CSE (10%) for 48 h were harvested and dual stained with AnnexinV and propidium iodide for flow cytometric analysis. Apoptotic cells are identified as being both AnnexinV and propidium iodide positive, incorporating both early and late stages of cell death. Data represents plots derived from one of two independent experiments.

**Suppl Figure 4. Nicotine treatment results in a modest and variable upregulation in ACE2 relative to CSE treatment (Donor 1)**. Nicotine treatment (1 µM) for 48 h also modestly increases ACE2 protein expression relative to untreated control. Calu-3 are presented as the positive control. Representative western blot from 3 independent experiments. WB performed using ACE2 N (Ab228349) or C-terminal (Ab15348) antibodies.

**Suppl Figure 5. CSE exposure does not significantly alter interferon-associated gene expression (Donor 1).** RT-PCR data shows log2 relative fold-change in expression from n=7 independent experiments from 7 experiments, (Mann-Whitney, ns).

**Suppl Figure 6.** CSE exposure does not promote an increase in dACE2 protein. Western blot showing 5 independent experiments showing the impact of CSE on flACE2/dACE2 expression. ACE2 antibody used ab15348.

**Suppl Figure 7. Impact of CSE and oltipraz on flACE2 expression and cellular infection**

**A.**  Representative western blots from two donors. Calu-3a and b represent Calu-3 cells from different experiments. ACE2 antibody (Ab15348). **B.** Quantitation of immunofluorescence signal using microscopy as described in Figure 5E - replicate experiment using cells at ALI on transwells from Donor 1 infected with 1x10^4^ TCID50 of B.1.1.7 lineage SARS-CoV-2. Antibody used was SARS-CoV-2 nucleocapsid MRC PPU DA114.

**Suppl Figure 8.** Figure based on RNA-Seq data from Gindele et al, 2020 extracted from Gene Expression Omnibus (GSE135188). Small airway epithelial cultures (SAEC) at ALI were exposed to cigarette smoke (CS) or air three times a week during the differentiation phase, that is from the period from Day 0 to Day 28. ACE2 expression is significantly increased by CS exposure in 2/6 donors whilst shows a positive trend for increasing in 4/6 donors.
