## Supplementary Figure Legends File for "Cigarette smoke preferentially induces full length ACE2 exposure in primary human airway cells but does not alter susceptibility to SARS-CoV-2 infection"

#### Suppl Figure 1

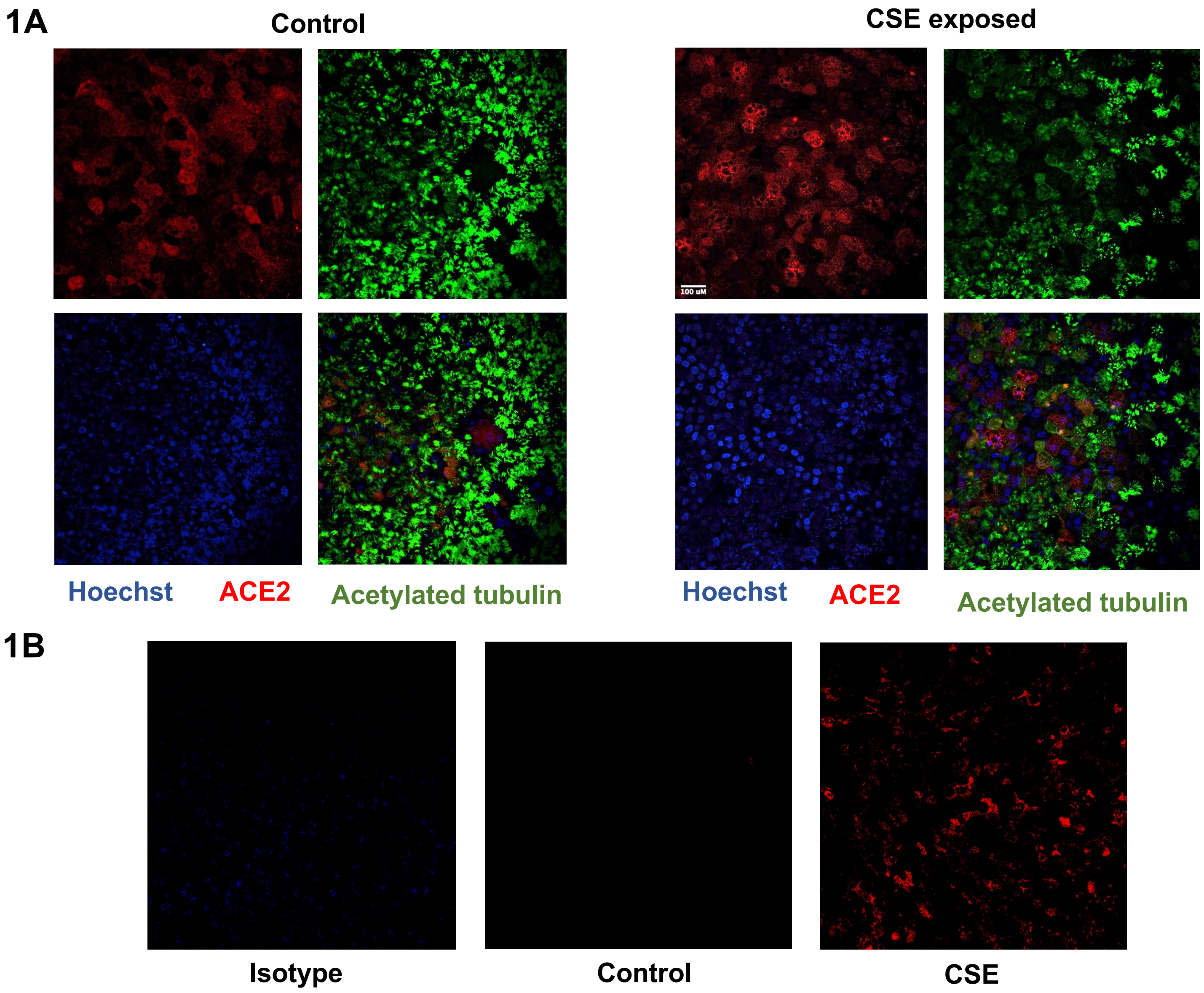

**Suppl Figure 2**

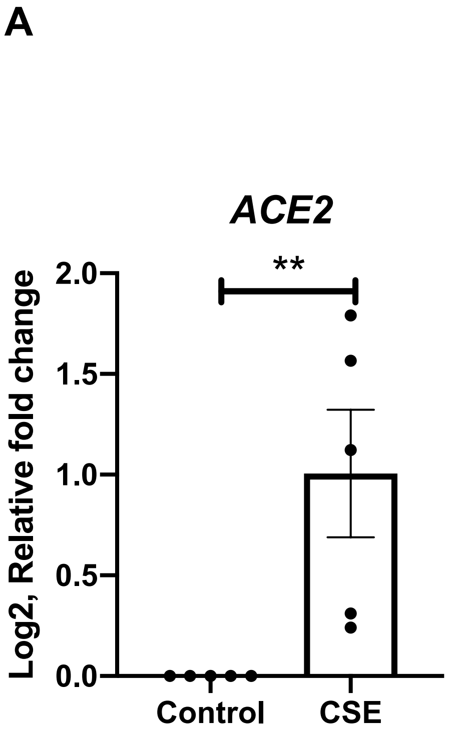


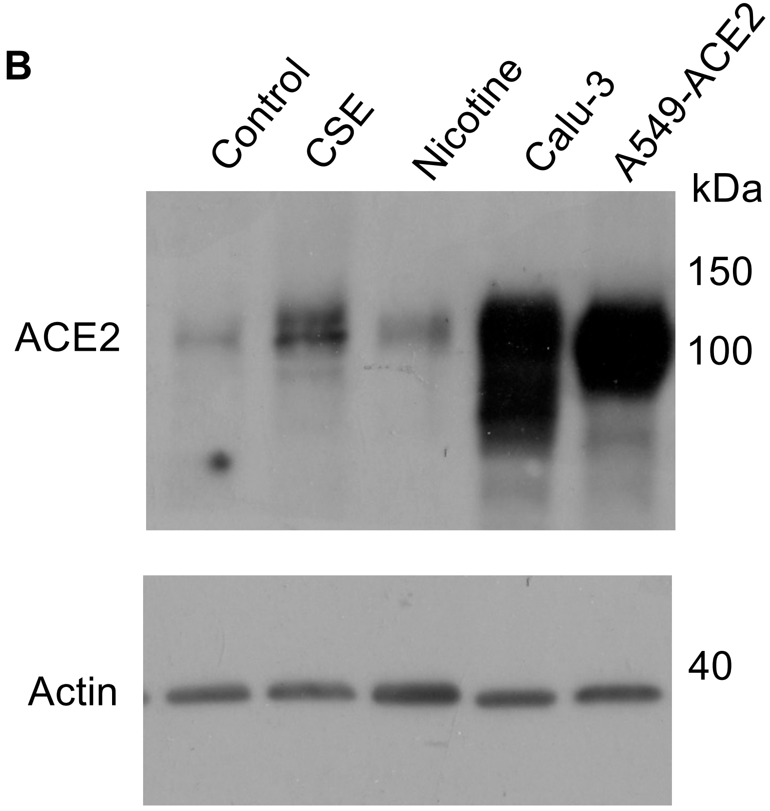

#### Suppl figure 3

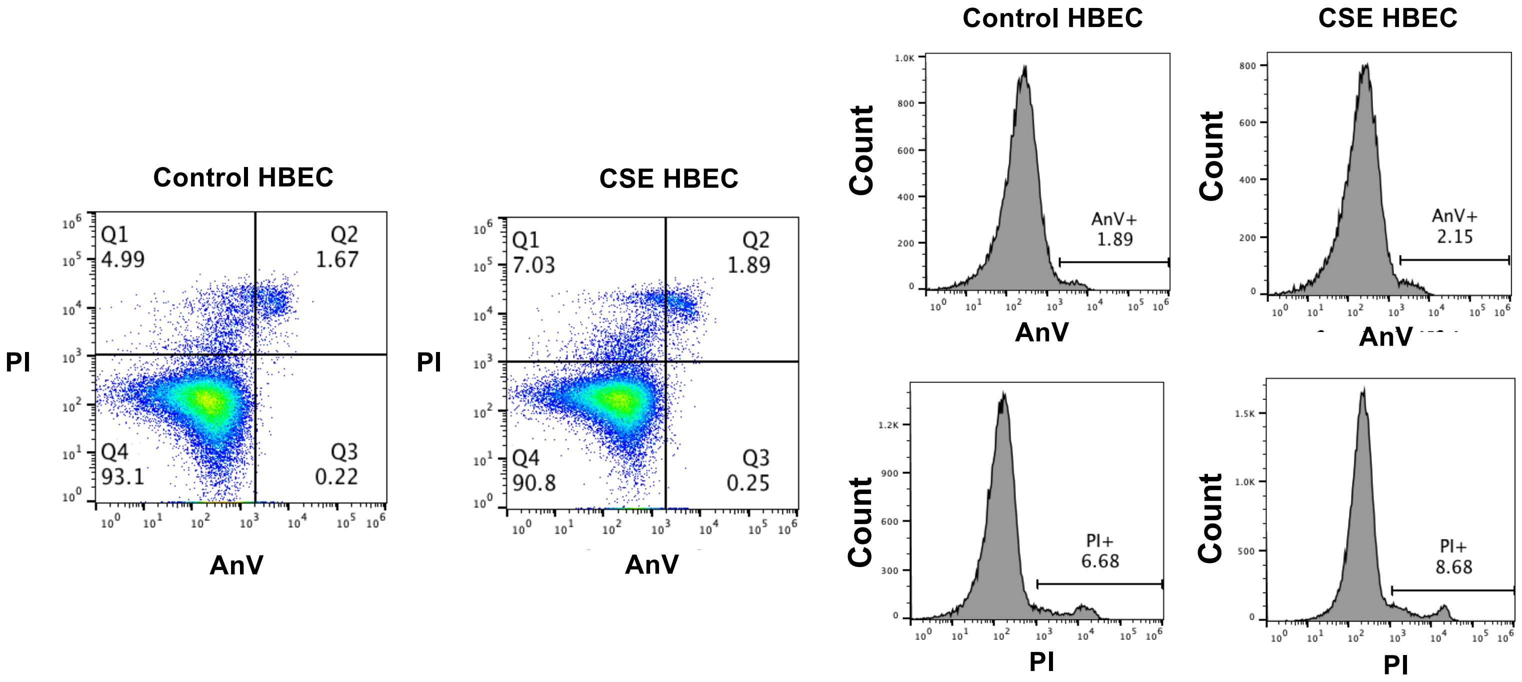

**Suppl figure 4**

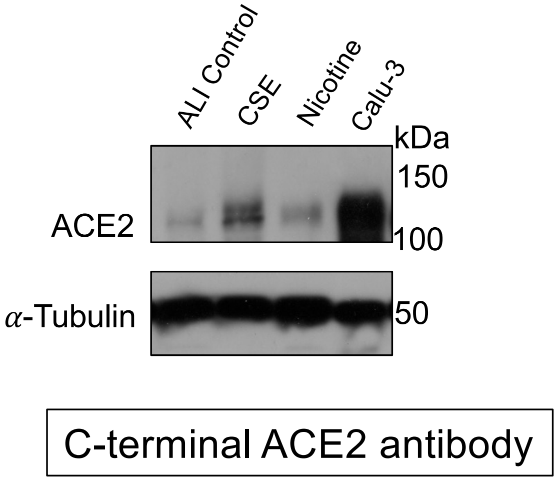


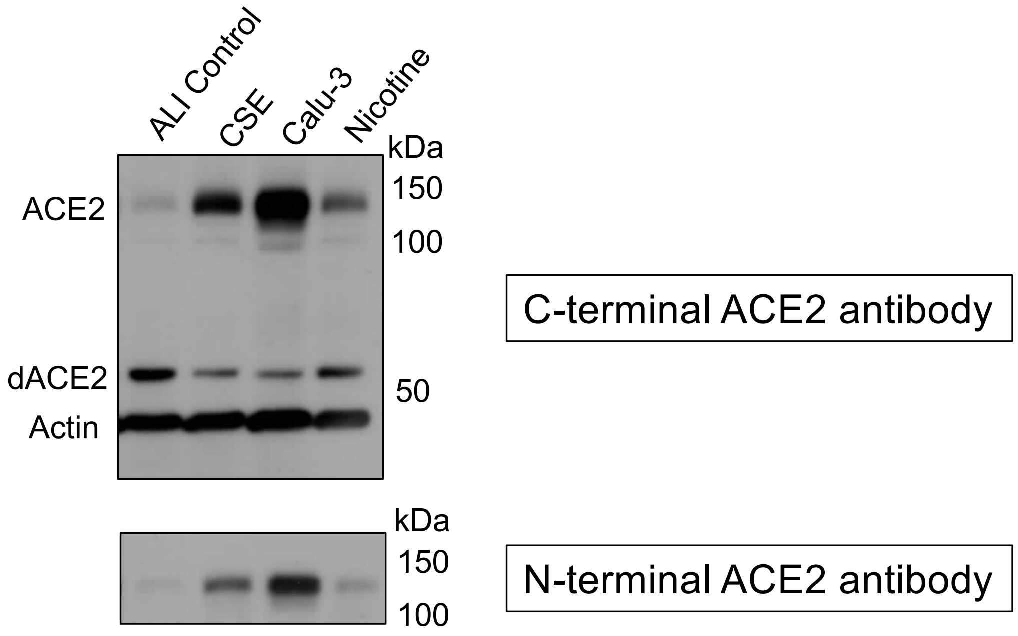

**Suppl figure 5**

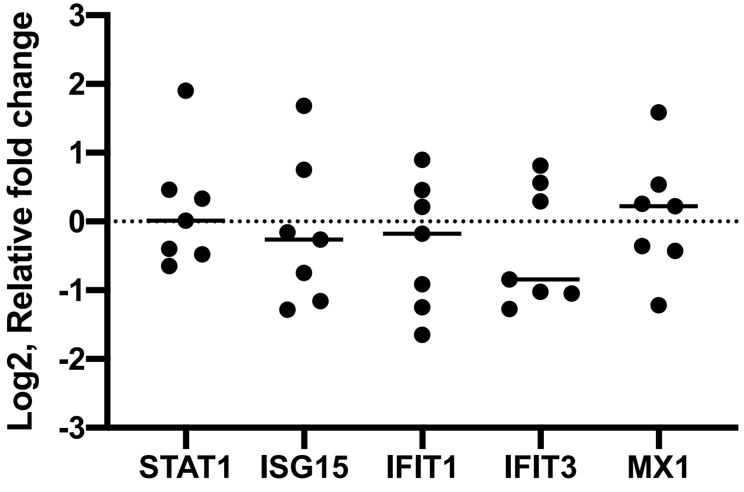

#### Suppl figure 6

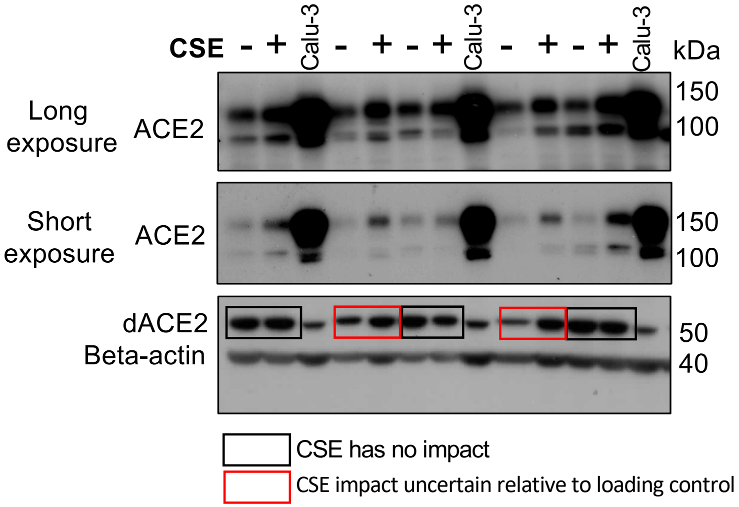

#### Suppl figure 7

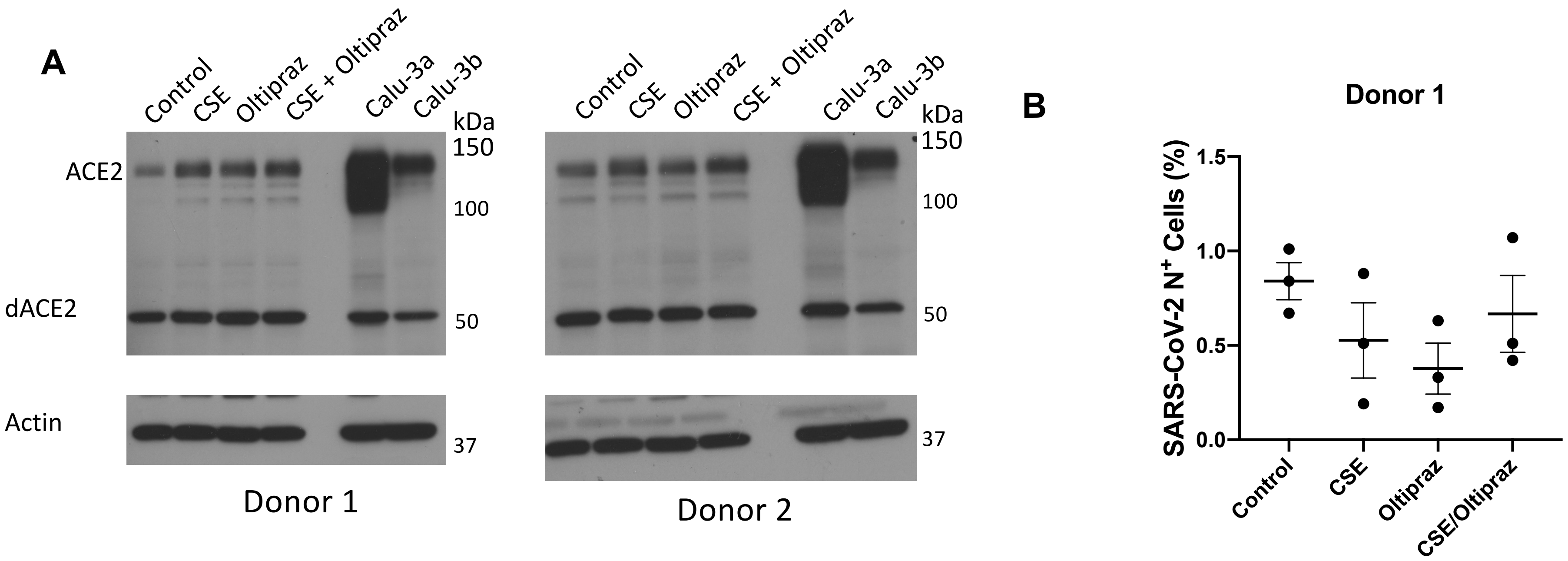

### Suppl figure 8

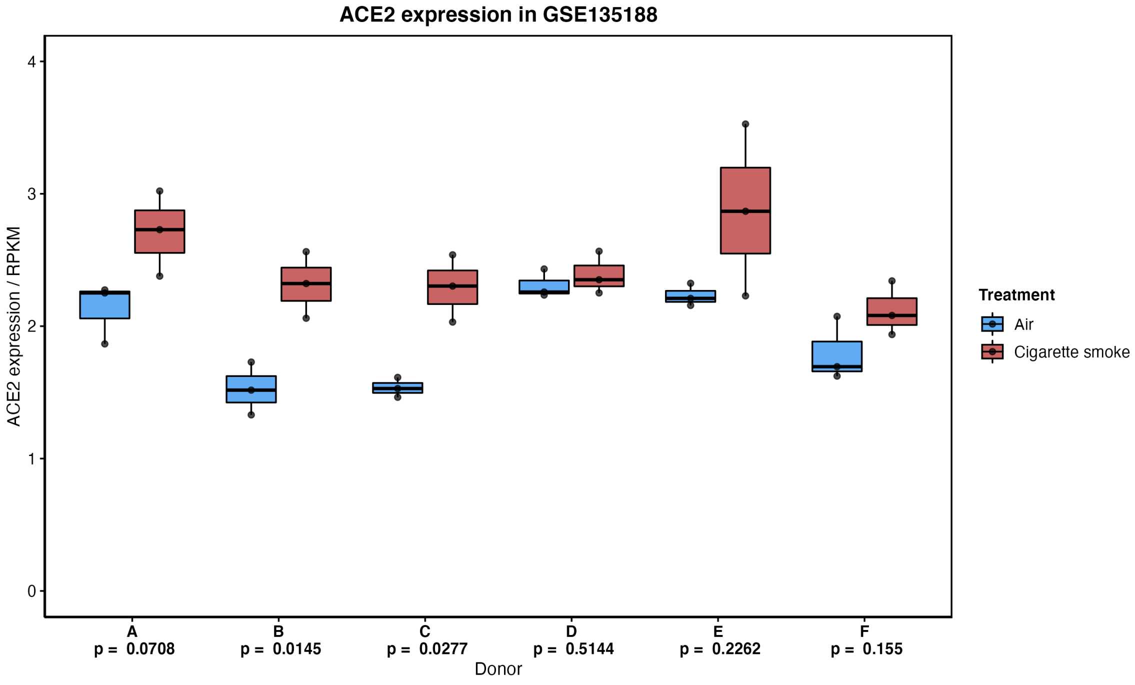
